# Kinetic and structural characterization of human NUDIX hydrolases NUDT15 and NUDT18 as catalysts of isoprene pyrophosphate hydrolysis

**DOI:** 10.1101/2023.11.29.569174

**Authors:** Emma R. Scaletti, Judith E. Unterlass, Ingrid Almlöf, Tobias Koolmeister, Karl S. Vallin, Despina Kapsitidou, Thomas Helleday, Pål Stenmark, Ann-Sofie Jemth

**Affiliations:** Department of Biochemistry and Biophysics, Stockholm University, 106 91 Stockholm, Sweden; Science for Life Laboratory, Department of Oncology-Pathology, Karolinska Institutet, 171 76 Stockholm, Sweden

**Keywords:** hydrolase, enzyme catalysis, enzyme kinetics, isoprenoid, protein structure, hydrolysis, isoprene pyrophosphate, NUDIX, NUDT15, NUDT18

## Abstract

Isoprene pyrophosphates play a crucial role in the synthesis of a diverse array of essential nonsterol and sterol biomolecules, and serve as substrates for post-translational isoprenylation of proteins, enabling specific anchoring to cellular membranes. Hydrolysis of isoprene pyrophosphates would be a means to modulate their levels, downstream products, and protein isoprenylation. While NUDIX hydrolases from plants have been reported to catalyze the hydrolysis of isoprene pyrophosphates, homologous enzymes with this function in animals have not yet been identified. In this study, we screened an extensive panel of human NUDIX hydrolases for activity in hydrolyzing isoprene pyrophosphates. We found that human NUDT15 and NUDT18 efficiently catalyze the hydrolysis of several physiologically relevant isoprene pyrophosphates. Notably, we demonstrate that geranyl pyrophosphate is an excellent substrate for NUDT18, which displays a catalytic efficiency of 2.1·10^5^ M^-1^s^-1^, thus making it the best substrate identified for NUDT18 to date. Similarly, geranyl pyrophosphate proved to be the best isoprene pyrophosphate substrate for NUDT15, with a catalytic efficiency of 4.0·10^4^ M^-1^s^-1^. LC-MS analysis of NUDT15 and NUDT18 catalyzed isoprene pyrophosphate hydrolysis revealed the generation of the corresponding monophosphates and inorganic phosphate. Furthermore, we solved the crystal structure of NUDT15 in complex with the hydrolysis product geranyl phosphate at a resolution of 1.70 Å. This structure revealed that the active site nicely accommodates the hydrophobic isoprenoid moiety and aided in identifying key binding residues. By overexpressing NUDT15 and NUDT18 in cells, we demonstrated a decrease in cellular cholesterol levels. Collectively, our findings strongly imply that isoprene pyrophosphates are endogenous substrates of NUDT15 and NUDT18, and support their involvement in animal isoprene pyrophosphate metabolism.

## INTRODUCTION

Isoprene pyrophosphates are metabolites in the mevalonate pathway which is an essential metabolic pathway found in eukaryotes, archaea, and some prokaryotes [1] and is responsible for the multistep conversion of acetyl-CoA into isopentenyl pyrophosphate (IPPP) and its isomer dimethylallyl pyrophosphate (DMAPP). IPPP and DMAPP are used as building blocks in the biosynthesis of all mammalian isoprenoids. In addition, IPPP and DMAPP are used to produce a diverse array of essential nonsterol and sterol biomolecules. An example is cholesterol which is crucial for maintaining cell membrane integrity and used for production of steroid hormones and bile acids. Other products derived from isoprene PPs are lipoproteins, vitamin K, heme A, the mitochondrial electron carrier ubiquinone, dolichol, and isopentenyladenine used in tRNA modification [1, 2]. Consequently, the isoprene PPs produced by the mevalonate pathway are crucial for a plethora of important biological processes. Figure 1 shows the structures of the physiologically relevant isoprene pyrophosphates (isoprene PPs) in animals.

**Figure 1.**
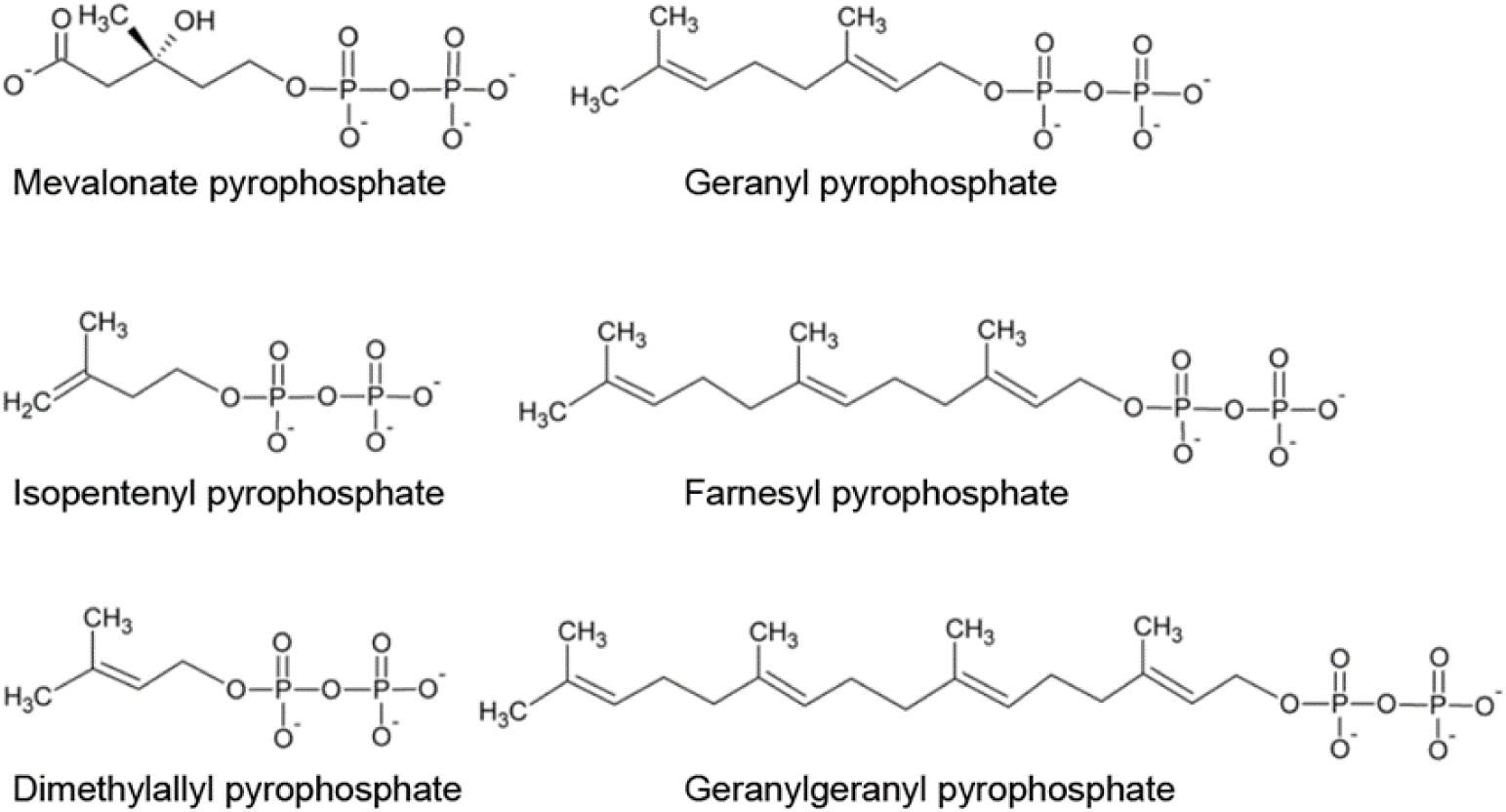
Structures of physiologically relevant isoprene-PPs.

Moreover, approximately 2% of all mammalian proteins undergo post-translational isoprenylation [3]. This process involves the covalent attachment of hydrophobic C-15 isoprene farnesyl or the C-20 isoprene geranylgeranyl group to cysteine residues in the C-terminal end of various proteins. Proteins that are farnesylated include the small GTP-binding proteins (GTPases) exemplified by the Ras superfamily, whereas other small GTPases such as Rab and Rho GTPases are geranylgeranylated [4]. Prenylation of these proteins enables their attachment to cellular membranes, a prerequisite for these proteins to exert their specific functions in biological processes such as cell survival, cell signaling, and proliferation [5].

The dysregulation of isoprenoid biosynthesis has been implicated in several biochemical disorders, underscoring the crucial role of appropriate isoprenoid levels in maintaining proper cellular function [6–8]. Additionally, statins, commonly employed to reduce cholesterol levels, exhibit effects beyond cholesterol regulation likely mediated by changes in isoprene metabolite levels [9]. These statin-induced alterations impact fundamental processes such as cell cycle regulation, cell migration, cell proliferation and cell survival [7, 10]. Given the significance of isoprenoid metabolites, their levels are presumably tightly controlled. Mevalonate kinase (MK), the second enzyme in the isoprene biosynthesis pathway, is subject to feedback inhibition by intermediates of the isoprenoid pathway - geranyl pyrophosphate (GPP), farnesyl pyrophosphate (FPP) and geranylgeranyl pyrophosphate (GGPP) [11]. Furthermore, HMG-CoA reductase, which catalyzes the rate-limiting step in sterol and isoprenoid biosynthesis, is regulated through a feedback loop to maintain cholesterol levels within appropriate ranges while nonsterol isoprenoids are still being produced. Sterol molecules and geranylgeraniol (GGOH) participate in this feedback loop by promoting the proteasomal breakdown of the HMG-CoA reductase enzyme [12]. In addition, geraniol (GOH) and farnesol (FOH) have been demonstrated to have inhibitory effects on HMG-CoA reductase activity [13–16]. How these isoprenoid alcohols are produced in human cells remains unclear, however, a membrane bound polyisoprenoid diphosphate phosphatase (PDP1) has been suggested to be involved in this process [17, 18]. However, no cytosolic phosphatases capable of catalyzing the hydrolysis of isoprene pyrophosphates have been identified to date. Building upon the finding that the NUDIX enzyme NUDTX1 in the plants *Arabidopsis thaliana* and rose (*Rosa hybrid cultivar*) can convert GPP to Geranyl monophosphate (GP) [19, 20], we hypothesized that enzymes within the human NUDIX enzyme family might possess the capability to hydrolyze isoprene pyrophosphates in the cell. To investigate this, we screened a panel of human NUDIX enzymes for activity with the isoprene pyrophosphates IPPP and DMAPP and found NUDT15 and NUDT18 to efficiently catalyze their hydrolysis. Moreover, we identified additional isoprene PPs as substrates of NUDT15 and NUDT18 of which GPP was found to be hydrolyzed with the highest catalytic efficiency. Notably, we showed that overexpression of NUDT15 and NUDT18 resulted in decreased cellular cholesterol content indicating that their activity with isoprene PPs plays a role in a cellular context. In summary, our findings unveil previously unknown hydrolysis activities of NUDT18 and NUDT15 with endogenous isoprene pyrophosphate substrates, suggesting a novel role for these enzymes in isoprene PP metabolism.

## RESULTS

### NUDT15 and NUDT18 are efficient catalysts of isoprene PP hydrolysis

To investigate whether any of the human NUDIX enzymes possess the capability to catalyze the hydrolysis of physiologically relevant isoprene PPs, we screened an extensive panel of 17 human NUDIX enzymes for activity with DMAPP and IPPP. Out of the enzymes tested, NUDT15 and NUDT18, exhibited activity with these isoprene PPs, as depicted in Figure 2A. Next, we tested the activities of NUDT15 and NUDT18 with additional isoprene-PPs and compared them with activities towards their previously described substrates, 8-oxo-dGDP for NUDT18 and dGTP for NUDT15. At a substrate concentration of 100 µM both NUDT15 and NUDT18 demonstrated significant hydrolysis activity with IPPP, DMAPP, GPP and FPP as illustrated in Figure 2B. Notably, only NUDT18 catalyzed the hydrolysis of mevalonate pyrophosphate (MPP). Furthermore, NUDT18 exhibited a considerably higher specific activity with GGPP compared to NUDT15. In addition, we assessed the activity of *E. coli* MutT, a bacterial homolog to NUDT1 and NUDT15, with GPP. However, no activity was detected (Figure S1), suggesting that the observed activity with isoprene pyrophosphates within the NUDIX family likely emerged at a later stage in evolution.

**Figure 2.**
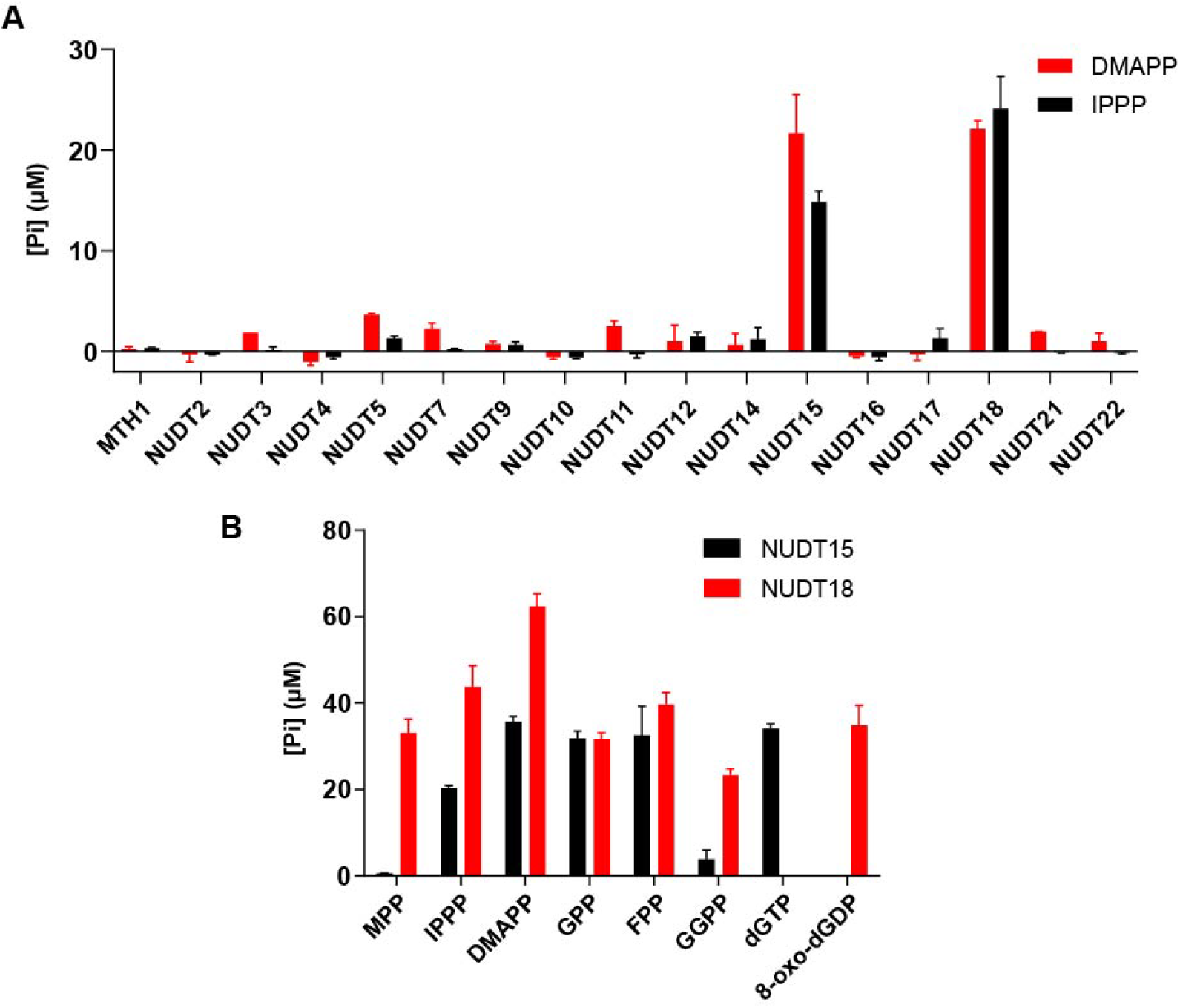
NUDT15 and NUDT18 are catalysts of isoprene pyrophosphate hydrolysis. (A) NUDT15 and NUDT18 catalyze the hydrolysis of the mevalonate pathway metabolites Dimethylallyl-PP and Isopentenyl-PP. A panel of 17 NUDIX hydrolases were screened for activity with Dimethylallyl-PP and Isopentenyl-PP using 100 nM human NUDIX enzyme and 100 µM substrate. (B) Activity of NUDT15 and NUDT18 (100 nM) was tested with isoprene pyrophosphate substrates (100 µM) and compared with activity towards the previously known substrates dGTP for NUDT15 and 8-oxo-dGDP for NUDT18.

### LC-MS analysis reveals hydrolysis takes place between phosphate groups

To further analyse the hydrolysis reactions of GPP and FPP, these isoprene PPs were incubated with NUDT15 and NUDT18, samples from the reaction mixtures were collected after different incubation times and subjected to LC-MS analysis. The detected mass (m/z=257 [M+Na^+^]) of the product peak was consistent with GP formation after GPP hydrolysis catalyzed by both NUDT18 (Figure 3A) and NUDT15 (Figure 4A), similar to the observations for AtNUDX1-catalyzed hydrolysis of GPP [20]. The time-dependent hydrolysis of GPP could be effectively tracked and was completed for NUDT15 and NUDT18 after 60 minutes and 90 minutes, respectively (Figure 3C and Figure 4C). The faster conversion of GPP in the case of NUDT15 is likely due to the two-fold higher concentration of NUDT15 compared to NUDT18 used in the assay. LC-MS analysis of FPP hydrolysis showed a product peak with m/z=325 [M+Na^+^], consistent with FP being the product after NUDT18 and NUDT15 catalyzed FPP hydrolysis (Figure 3B and Figure 4B). In comparison to the hydrolysis of GPP, the hydrolysis of FPP is notably slower, especially for NUDT15, where hydrolysis appears to plateau at 50% conversion. This may potentially be due to product inhibition of the enzyme or be due to low solubility of the hydrophobic FP. Additionally, in Figure 3A an extra small peak is visible just below 1.6 minutes elution time after NUDT18 catalyzed hydrolysis of GPP for 180 minutes. This peak may represent GOH that is formed by NUDT18 catalyzed hydrolysis of the produced GP.

**Figure 3.**
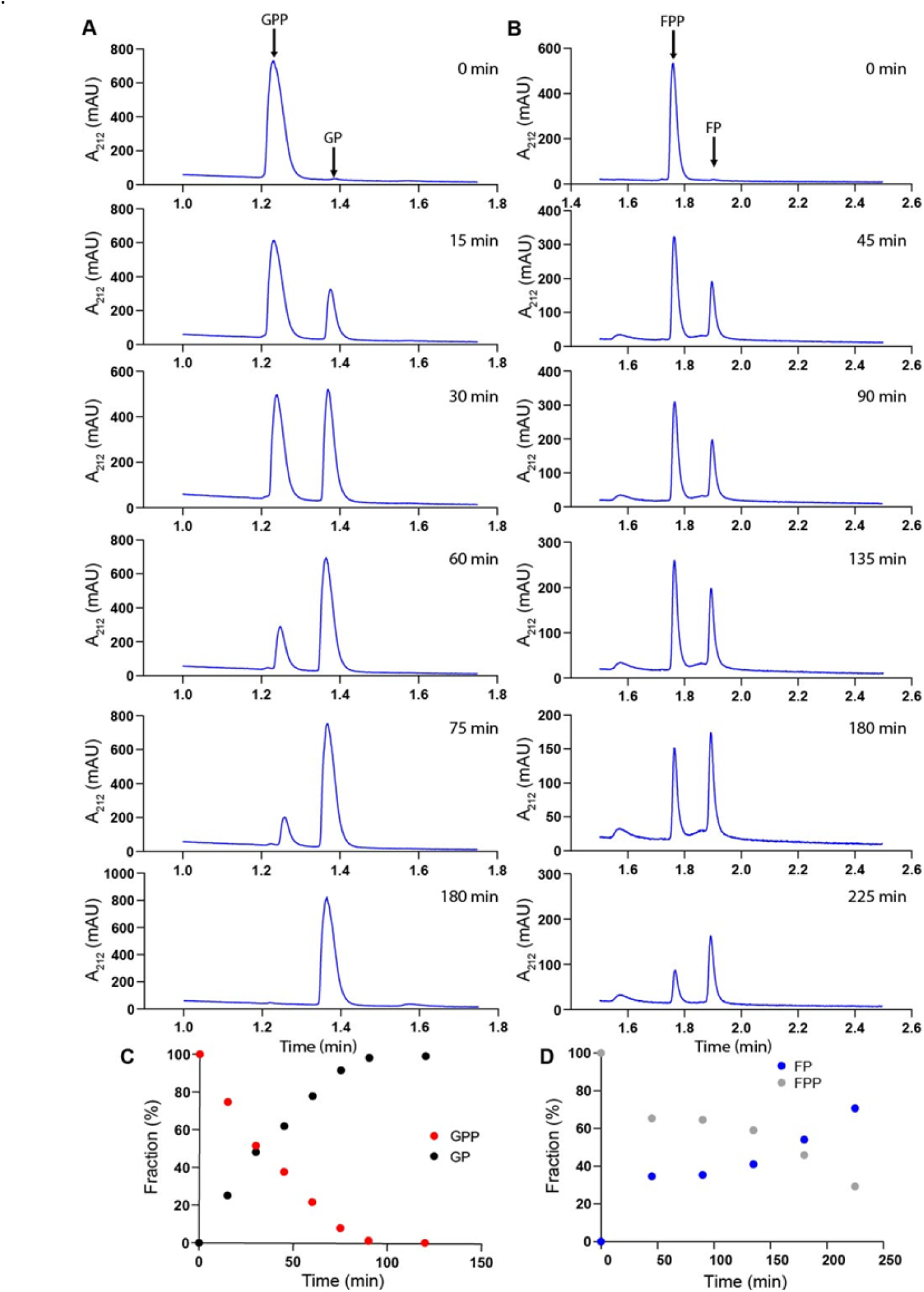
LC-MS analysis of NUDT18 catalyzed hydrolysis of GPP and FPP reveals isoprene monophosphates production. HPLC analysis showing that hydrolysis takes place between the phosphate groups in GPP and FPP and produces GP and FP. NUDT18 catalyzed hydrolysis of GPP (A) and FPP (B) was followed over time by incubation of GPP or FPP with NUDT18 and analysis of reaction mixtures at different time points using HPLC and MS. The fraction of hydrolyzed substrate and formed product was calculated for GPP hydrolysis (C) and FPP hydrolysis (D) using the area under the curve of the peaks.

**Figure 4.**
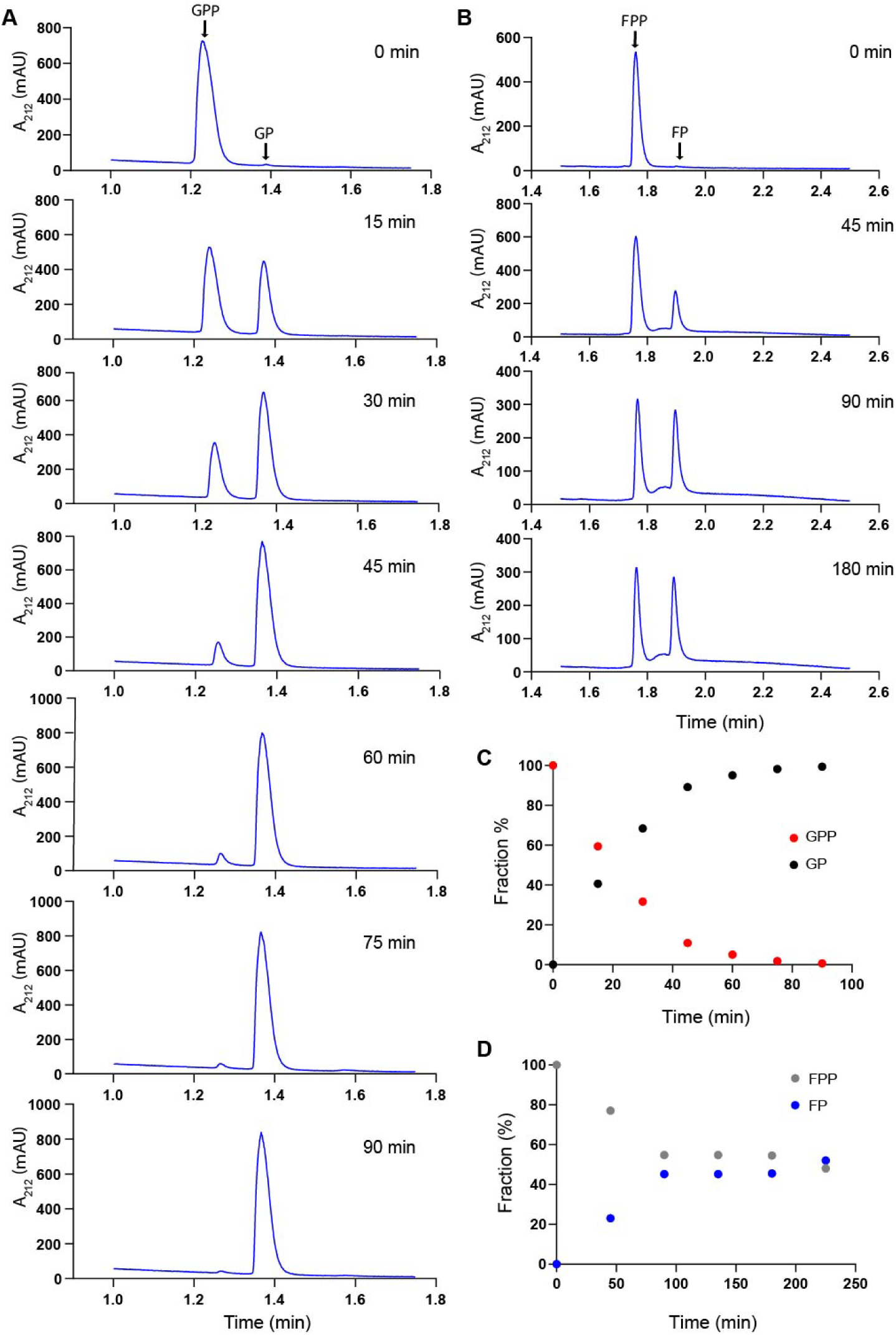
LC-MS analysis of NUDT15 catalyzed hydrolysis of GPP and FPP reveals production of isoprene monophosphates. HPLC analysis showing that hydrolysis takes place between the phosphate groups and produces GP and FP. NUDT15 catalyzed hydrolysis of GPP (A) and FPP (B) was followed over time by incubation with NUDT15 and sampling at different time points. Samples were analysed using HPLC and MS and the fraction of hydrolyzed substrate and formed product was calculated for GPP hydrolysis (C) and FPP hydrolysis (D) using the area under the curve of the corresponding peaks.

### Detailed kinetic analysis of NUDT15 and NUDT18 with isoprene PPs

To further investigate the kinetics of NUDT18 and NUDT15 catalyzed hydrolysis of the identified isoprene PP substrates, we produced saturation curves by measuring initial enzyme activity rates at 22 °C at a range of substrate concentrations (Figure 5A, B, and C). The decision to assay the reaction rates at 22°C rather than the more physiologically relevant temperature of 37°C was based on the excessively rapid consumption of substrates at the lower concentrations, which hindered the accurate assessment of initial rates at 37 °C. Determined kinetic parameters for NUDT18 and NUDT15 are presented in Table 1 and Table 2, respectively. NUDT18 displays comparable turnover numbers for GPP, DMAPP and IPPP whereas the corresponding values for MPP and FPP are slightly lower. The Km-value of NUDT18 was found to be lowest for GPP (3.7 µM) indicating that the affinity is highest for this substrate, followed by DMAPP, IPPP, FPP, GGPP and MPP. The catalytic efficiency (kcat/Km) of NUDT18 is highest for GPP followed by DMAPP, IPPP, FPP, MPP and GGPP. However, the calculated low catalytic efficiency of NUDT18 with GGPP might be underestimated due to the limited solubility of this highly hydrophobic substrate (Figure 1) in the aqueous assay buffer used. It is notable that the catalytic efficiency of NUDT18 catalyzed hydrolysis of GPP is considerably higher than what has been determined for the previously reported endogenous substrate, 8-oxo-dGDP [21].

**Figure 5.**
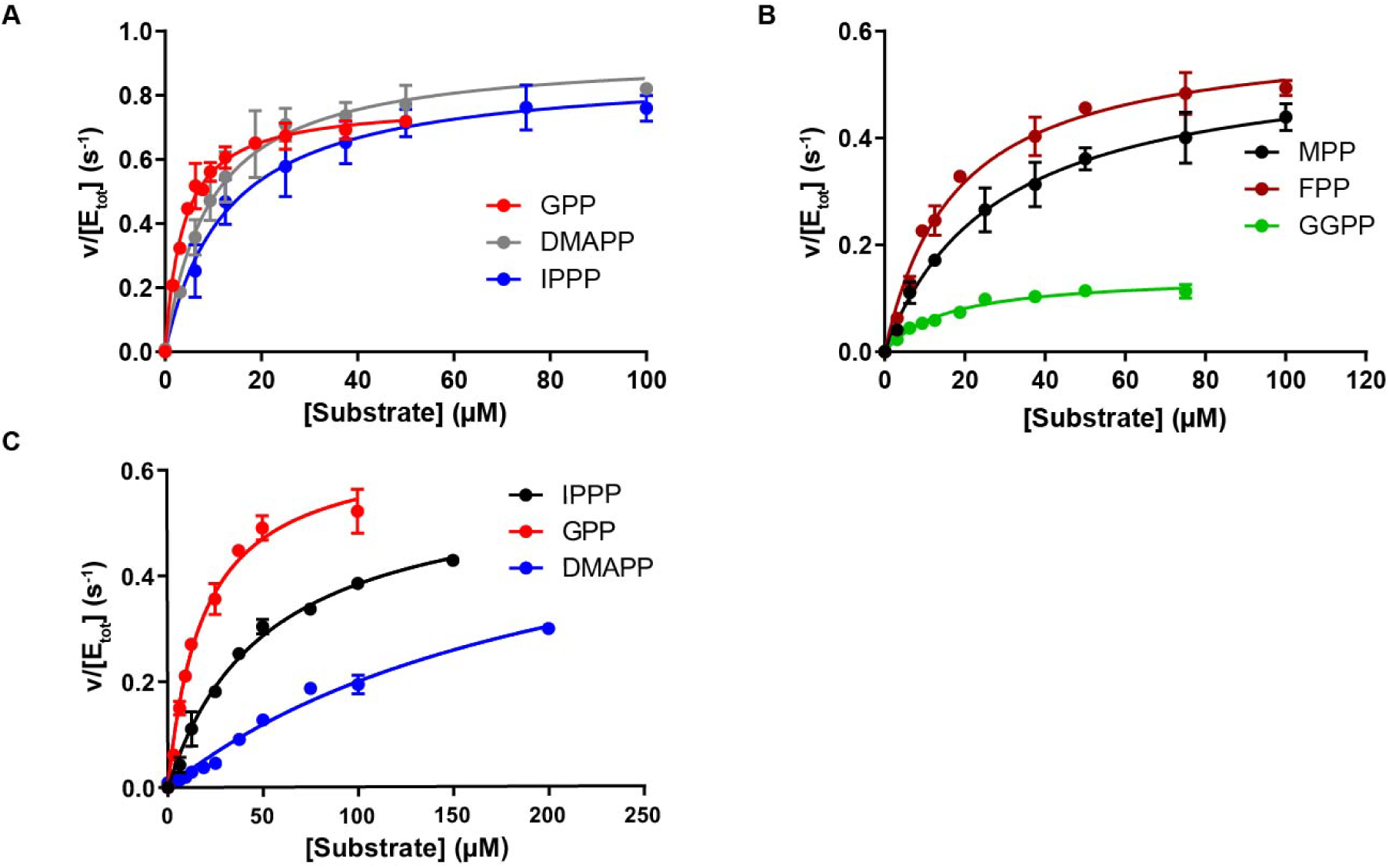
NUDT18 and NUDT15 are efficient catalysts of GPP hydrolysis. (A) Saturation curves of NUDT18 with GPP, DMAPP and IPPP. (B) Saturation curves of NUDT18 with MPP, FPP and GGPP. (C) Saturation curves of NUDT15 with GPP, DMAPP and IPPP. Graphs show saturation curves with data points representing average and standard deviation from two independent experiments in which initial rates were determined using enzyme activities assayed in duplicate at three different time points.

**Table 1.** Kinetic parameters for NUDT18 with isoprene pyrophosphates.

| Kinetic parameter |  | Substrate |  |  |  |  |  |
| --- | --- | --- | --- | --- | --- | --- | --- |
|  |  | MPP | IPPP | DMAPP | GPP | FPP | GGPP |
|  | n | 2 | 2 | 2 | 2 | 2 | 2 |
| $k_{\text{cat}}/K_m$ ($\text{M}^{-1}\text{s}^{-1}$) | Mean | 20,400 | 72,100 | 99,700 | 214,300 | 27,500 | 8,500 |
|  | SD | 400 | 19,800 | 27,500 | 25,900 | 900 | 1,200 |
| $K_m$ ($\mu\text{M}$) | Mean | 26.8 | 12.4 | 9.8 | 3.7 | 18.8 | 17.9 |
|  | SD | 2.8 | 2.6 | 2.9 | 0.4 | 3.9 | 3.4 |
| $k_{\text{cat}}$ ($\text{s}^{-1}$) | Mean | 0.55 | 0.87 | 0.94 | 0.78 | 0.52 | 0.15 |
|  | SD | 0.07 | 0.05 | 0.01 | 0.02 | 0.12 | 0.01 |

n, number of independent experiments, SD, standard deviation

**Table 2.** Kinetic parameters for NUDT15 with isoprene pyrophosphates.

| Kinetic parameter |  | Substrate |  |  |  |  |
| --- | --- | --- | --- | --- | --- | --- |
|  |  | GPP | IPPP | DMAPP | FPP | GGPP |
|  | n | 2 | 2 | 2 | 1 | 1 |
| $k_{\text{cat}}/K_m$ ($\text{M}^{-1}\text{s}^{-1}$) | Mean | 39,860 | 8,330 | 6,470 | 1,350 | 870 |
|  | SD | 7,160 | 4,370 | 4,770 |  |  |
| $K_m$ ($\mu\text{M}$) | Mean | 18.7 | 63.6 | 159.3 | 76.3 | 163.3 |
|  | SD | 0.2 | 19.4 | 40.5 |  |  |
| $k_{\text{cat}}$ ($\text{s}^{-1}$) | Mean | 0.72 | 0.49 | 0.94 | 0.10 | 0.10 |
|  | SD | 0.11 | 0.11 | 0.50 |  |  |

n, number of independent experiments, SD, standard deviation

GPP was also the best substrate for NUDT15 among the isoprene pyrophosphates tested (Figure 5C and Table 2). NUDT15 displays both the highest kcat-value (0.7 s^-1^) as well as the lowest Km-value (18.7 µM) with GPP resulting in a catalytic efficiency of 39,900 M^-1^s^-1^. However, this is more than 5-fold lower compared to the corresponding catalytic efficiency of NUDT18 with GPP. The second best substrate for NUDT15 is IPPP followed by DMAPP with catalytic efficiencies of 8300 M^-1^s^-1^ and 6500 M^-1^s^-1^, respectively.

### Co-crystallization and structure determination of the NUDT15-GP complex

In order to understand the binding mode of the isoprene PPs we co-crystalized NUDT15 with GPP. Extensive efforts were also made to produce crystals of NUDT18 with GPP and other isoprene PPs but without success. We solved the structure of human NUDT15 in complex with GP to 1.70 Å resolution. The tertiary structure of NUDT15 is a homodimer, consistent with biochemical studies that show the protein is also dimeric in solution [22]. The NUDT15 monomer is comprised of two α-helices, seven β strands and three 310-helices (Figure 6A).

**Figure 6.**
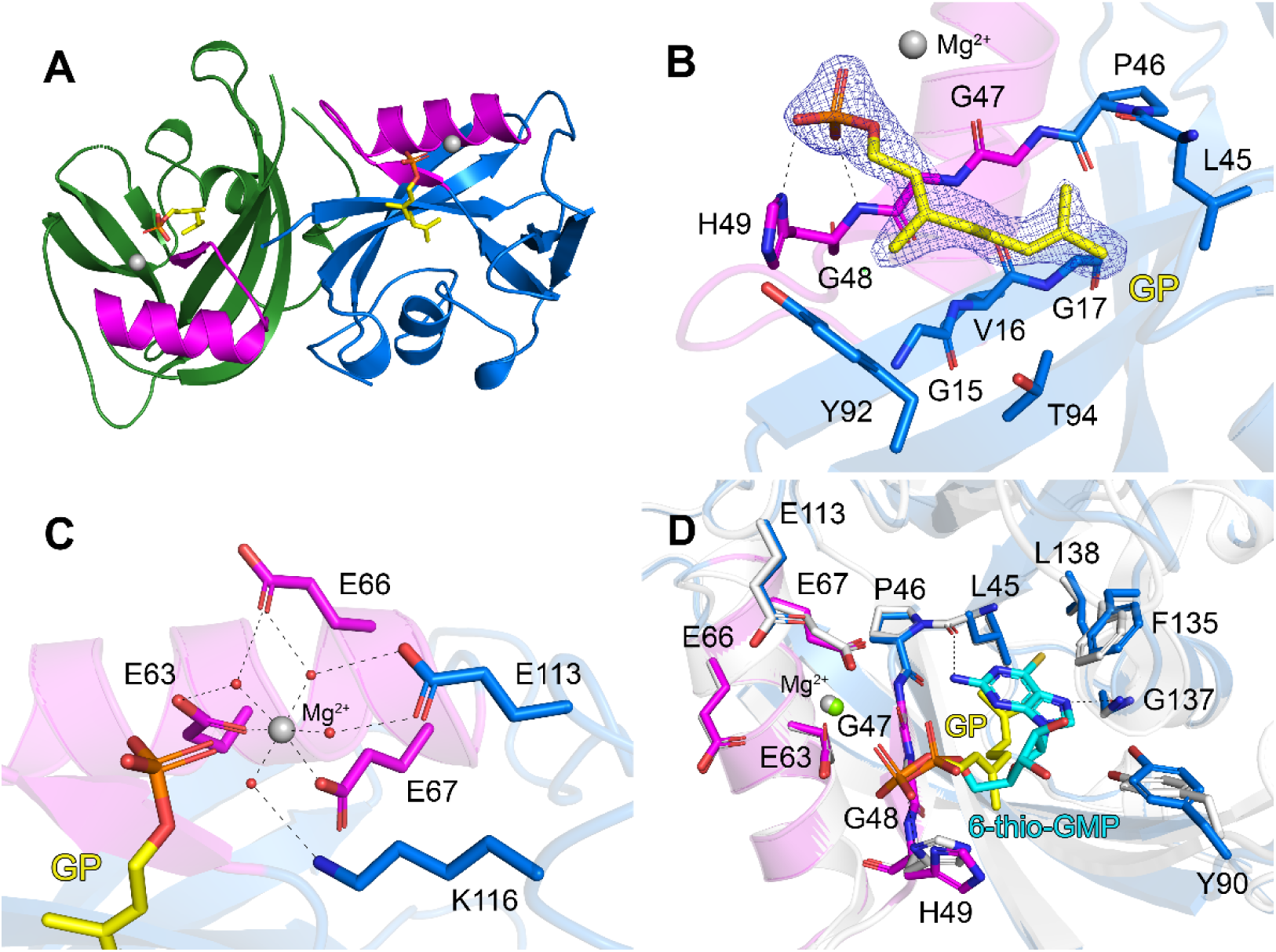
Crystal structure of NUDT15 in complex with GP. (A) Overall structure of the NUDT15 dimer, depicted as a ribbon representation. Each monomer is colored green and blue, respectively. The highly conserved NUDIX motif is colored magenta. The product GP is shown as a stick representation with its respective atoms colored; carbons yellow, oxygens red and phosphors orange. The magnesium ions required for activity and ligand coordination are shown as grey spheres. (B) Active site hydrogen bond network of GP. Residues contributing to ligand binding are labeled by residue number. Nitrogen and oxygen atoms are colored blue and red, respectively. Hydrogen bond interactions are shown as dashed lines. The calculated 2*F*o-*F*c electron density map around GP is shown in blue, contoured at 1.0 σ and the *F*o-*F*c electron density maps are green and red contoured at +3.5 σ and -3.5 σ, respectively. (C) Hydrogen bond network showing magnesium coordination in the active site. Water molecules are shown as red spheres. (D) C⍺-atom superposition of hNUDT15-GP (coloring identical to panels A-C) with hNUDT15-6-thio-GMP (colored white; PDB ID: 5lpg). The hydrogen bond network for 6-thio-GMP is shown and the bound magnesium ion is shown as a green sphere. 6-thio-GMP is shown as a stick model with its atoms colored; carbons cyan, nitrogens blue, oxygens red, phosphors orange and sulphurs gold.

The individual monomers of NUDT15 are virtually identical as indicated by the low rmsd value of 0.37 Å for the corresponding Cα-atoms. The conserved NUDIX family motif (GX5EX7REVXEEXGU), which contains the residues required for magnesium binding and substrate hydrolysis, is located on α-helix 1 and β-strand 4. After structure refinement, it was evident that each monomer had electron density consistent with a single GP, indicating efficient hydrolysis of the GPP substrate by NUDT15 during co-crystallization (Figure 6A). The GP product is nicely accommodated within the active site and is positioned by two hydrogen bonds between the alpha-phosphate and His49, along with extensive hydrophobic interactions with Gly15, Val16, Gly17, Leu45, Pro46, Gly47, Gly48, Try92 and Thr94 (Figure 6B). Each active site also contains a single magnesium ion exhibiting ideal octahedral coordination with the catalytic residues Glu63, Glu66 and four ordered water molecules, three of which are additionally supported by the residues Glu67, Glu113 and Lys116 (Figure 6C). Comparison of hNUDT15-GP with the structure of hNUDT15 in complex with 6-thio-GMP shows the structures superimpose well, with a low rmsd value of 0.47 Å. 6-thio-GMP occupies a deeper position in the binding pocket compared to GP and is coordinated by more extensive hydrogen bonds, including the sidechain of His49, the main-chain nitrogen of Gly137 and the main-chain oxygen of Leu45. 6-thio-GMP is also supported by numerous hydrophobic interactions involving Pro46, Gly47, Gly48, Try90, Phe135 and Leu138. Importantly, the alpha-phosphates of both GP and 6-thio-GMP occupy a similar position, as do the magnesium ions in each structure (Figure 6D).

### Comparison of hNUDT15-GP and hNUDT18 structures

In order to identify structural features that are important for the activity of NUDIX enzymes with isoprene PPs we superimposed the structures of hNUDT15-GP and the hNUDT18 nucleotidase domain (PDB ID: 3GG6). This showed that while the core structures overlay adequately, there are some regions that do not superimpose particularly well (rmsd for all C-alpha atoms of 2.44 Å, amino acid sequence identity 29 %) (Figure S2A). This is in part due to small conformational changes that occur upon substrate binding, which are not observed in the ligand and metal free NUDT18 structure. Analysis of the ligand binding pocket shows there is sufficient space for isoprenoid substrates in the NUDT18 active site (Figure S2B). Many of the hydrophobic interactions involved in GP binding and residues required for metal binding are conserved between the structures (Figure S2B and C). However, there are also notable differences between hNUDT15 and hNUDT18. Interestingly, it is evident that the GPP binding pocket of NUDT18 has a greater abundance of hydrophobic residues than the NUDT15 structure. Specifically, residues Gly15, Gly17 and Gly47 in NUDT15 correspond to valine, leucine and alanine in NUDT18 (Figure S2B and C). This presumably would give NUDT18 a higher affinity for GPP compared to NUDT15, explaining its superior activity towards this substrate. Two other important differences include His49 and Thr92, which are both arginine residues (Arg77 and Arg117) in NUDT18. As shown in our hNUDT15-GP structure, His49 hydrogen bonds with the phosphate group of GP through nitrogen atoms in both its main chain and sidechain, suggesting that an arginine at this position should also be able to perform this function. Arg117 from NUDT18 is particularly interesting in the context of substrate specificity towards MPP. Unlike the other isoprenoids we examined which have hydrophobic core structures, MPP contains a carboxyl group (Figure 1) that would not be favored in the hydrophobic binding pocket. Based on our superpositions, only NUDT18 has a residue (Arg117) within hydrogen bonding range of MPP. This may be one reason why NUDT18 is capable of hydrolyzing MPP, whereas NUDT15 is unable to do so.

### Comparison of hNUDT15-GP and AtNUDT1-IPP structures

NUDT1 from *Arabidopsis thaliana* has been reported to exhibit activity with isoprene PPs [20]. Comparison of the structures of hNUDT15-GP and AtNUDT1-IPP (PDB ID: 6DBZ) [23] reveals that the core structures superimpose quite well as indicated by the low rmsd value of 1.34 Å for their C-alpha atoms (amino acid sequence identity 42 %) (Figure S3A). Although a structure of AtNUDT1 in complex with GPP has been determined (PDB ID, 5GP0), it lacks magnesium ions necessary for proper substrate coordination [20]. We therefore focused our comparisons on an available AtNUDT1 isopentenyl phosphate complex, where the substrate is coordinated by three magnesium ions [23]. Interestingly, the hydrophobic portions of both IPP and GP occupy slightly different positions in the structures, as are their alpha-phosphates. In fact, the alpha-phosphate of GP superimposes more closely with the beta-phosphate of IPP (Figure S3B). Residues involved in magnesium coordination are completely conserved between the structures, and the single magnesium in hNUDT15-GP occupies the same position as the second magnesium in the AtNUDT1-IPP structure (Figure S3A and B). This positioning may provide the enzyme with the flexibility to accommodate molecules with different numbers of phosphate groups, enabling NUDT15 to efficiently hydrolyze both di-and triphosphate substrates. Upon comparing the GP binding sites, we observed that Gly15 and Gly17 in hNUDT15 corresponds to alanine and valine in AtNUDT1 (Figure S3C). In the AtNUDT1-IPP complex, the ligand forms hydrogen bonds with His42, Gly40 and Glu60, which are entirely conserved in hNUDT15. The substrate is also positioned by hydrophobic interactions involving Ala11, Tyr87 and Phe127. These residues are conserved with hNUDT15 except for Ala11 which is a glycine residue in the human enzyme (Figure S3C). Notably, AtNUDT1, like NUDT18, has a more hydrophobic binding pocket compared to NUDT15, potentially explaining why these enzymes hydrolyze isoprene PPs much more efficiently than NUDT15 [23] as the binding of the hydrophobic portion of the isoprenoid substrates rely on hydrophobic interactions. Clearly, even small differences here can result in large differences in terms of binding affinity and catalytic efficiency towards the isoprene-PP substrates.

In order to identify residues in NUDT15 that have been conserved during evolution, we performed a multiple sequence alignment of NUDT15 proteins from humans and a selection of other vertebrates (Figure S4). His49, the only active site residue forming a hydrogen bond in the hNUDT15-GP structure, is entirely conserved. Residues that coordinate magnesium are also completely conserved across the sequences with a single exception of Glu66, which is a glutamate in all sequences except for chicken NUDT15. The residues involved in hydrophobic interactions in the NUDT15-GP complex are generally conserved, except for Gly15, Val16 and Gly17, which are replaced by larger hydrophobic residues in some of the other sequences. This observation aligns well with what we observed in the hNUDT18 and AtNUDT1 structures, suggesting that other vertebrate NUDT15 enzymes should also be capable of hydrolyzing isoprene PPs. In addition, we performed a multiple sequence alignment of NUDT18 proteins from humans and a range of other vertebrates (Figure S5). All residues predicted to interact with the isoprenoids are fully conserved, indicating that NUDT18 enzymes from other vertebrate species would likely exhibit activity with isoprene pyrophosphates as well.

### His49 is crucial for GPP hydrolysis activity

Based on our structural comparisons of hNUDT15-GP with existing NUDT15 and AtNUDT1 complexes, it is evident that residue His49 plays a crucial role in positioning the phosphate group of the substrate, as well as stabilizing the monophosphate product. To investigate the impact of this residue on catalytic activity and substrate binding, we generated the NUDT15H49A mutant and determined the activities of NUDT15wt and NUDT15H49A with dGTP and GPP (Figure 7A). At presumably saturating concentrations of the two substrates (100 µM) the H49A mutation causes a dramatic decrease in activity for both substrates. To investigate the role of the H49A mutation in more detail we produced saturation curves for NUDT15wt and NUDT15H49A with GPP (Figure 7B) and dGTP (Figure 7C) and determined kinetic parameters (Table S2). The H49A mutation led to a 30-fold reduction of the kcat-value for GPP while causing only a 2-fold drop in the kcat-value for dGTP. However, the introduction of the H49A mutation resulted in a 14-fold increase in the Km-value for dGTP leading to a 28-fold reduction in catalytic efficiency. Determining an exact Km-value for GPP is not possible due to the very low activity of NUDT15H49A with this substrate. It appears that the His49A mutation has a more pronounced negative impact on the kcat-value of NUDT15 catalyzed hydrolysis of GPP compared to dGTP, likely because the phosphate groups of nucleotide phosphates have more extensive hydrogen bond interactions in the active site than the phosphate groups of isoprene diphosphate substrates, making the His49A mutation relatively less detrimental [23, 24]. Overall, it is evident that the H49A mutation affects the catalytic activity of NUDT15 with both GPP and dGTP.

**Figure 7.**
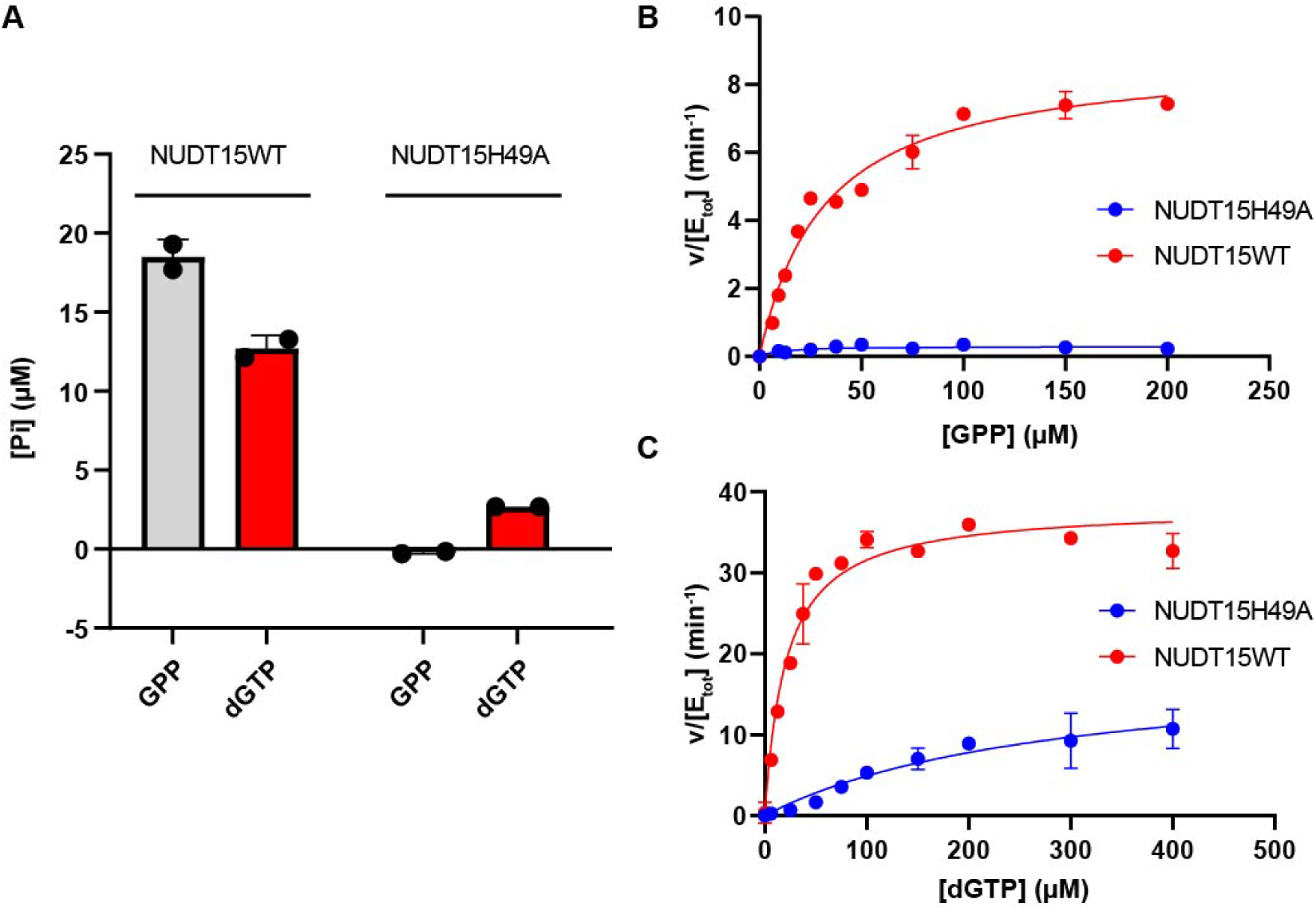
The H49A mutation drastically decreases hydrolysis activity with dGTP and GPP. (A) Activity was tested using 100 µM dGTP (red bars) or GPP (grey bars) and 100 nM NUDT15WT or NUDT15H49A at 22 °C. (B) Saturation curves of NUDT15WT and NUDT15H49A were produced by determining initial rates using 50 nM NUDT15 WT and 100 nM NUDT15H49A and substrate concentrations of GPP ranging from 0 to 200 µM. (C) Saturation curves with dGTP were produced by determining initial rates of 5 nM NUDT15WT or 50 nM NUDT15H49A at dGTP concentrations between 0 and 400 µM.

### NUDT15 and NUDT18 overexpression decreases cellular cholesterol

Based on the observed activity of NUDT15 and NUDT18 with isoprene PPs we hypothesized that this would have an impact on intracellular levels of isoprene PPs and consequently affect cellular cholesterol levels. To investigate this, we produced cell lines that overexpress NUDT15 and NUDT18 (Figure 8A and Figure S6A and B). NUDT15 and NUDT18 overexpressing cells and control cells carrying empty expression vectors were statin treated or left untreated (Figure 8B) and cholesterol levels were analyzed (Figure 8C, D and Figure S6).

**Figure 8.**
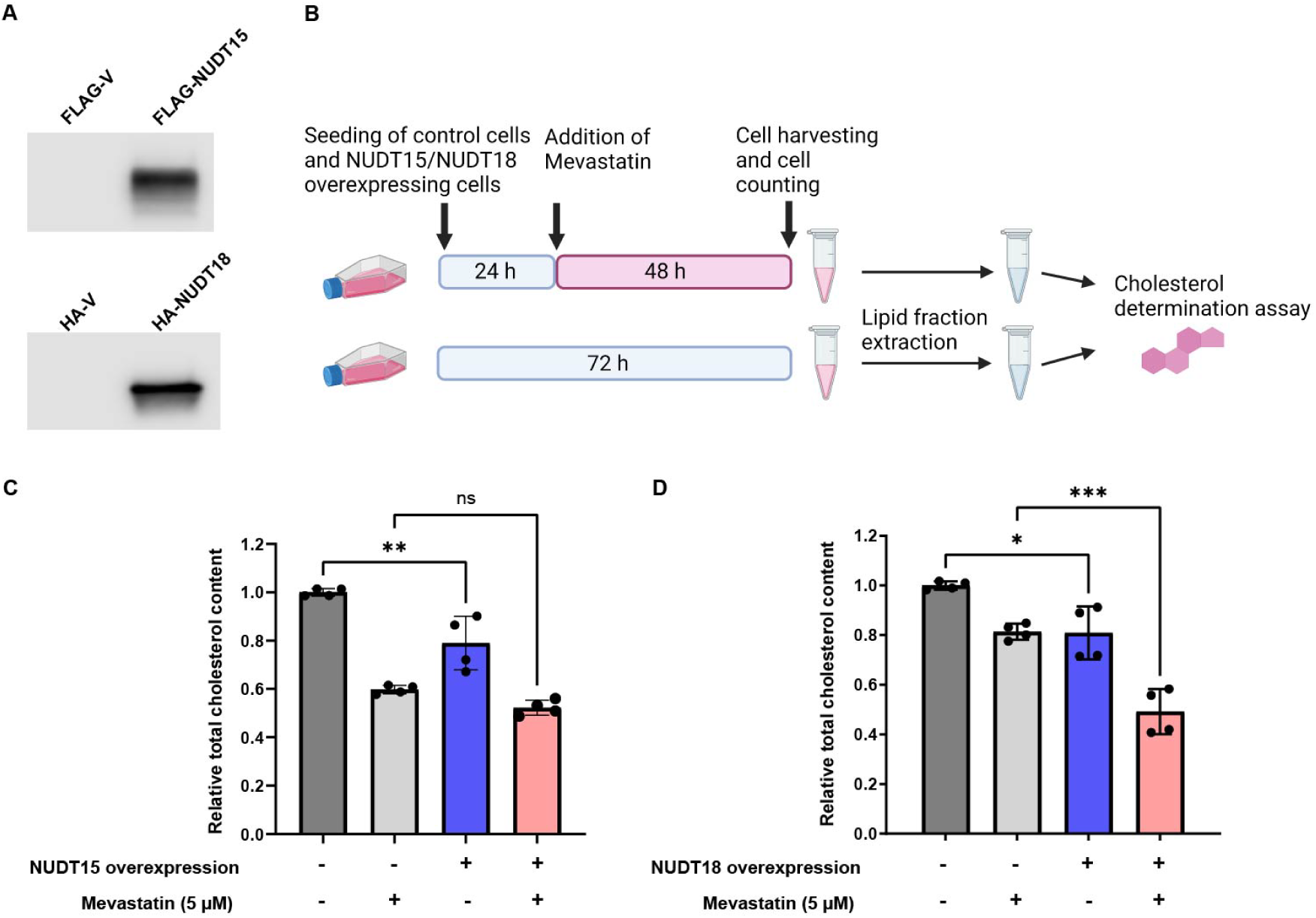
Overexpression of NUDT15 and NUDT18 affects cellular cholesterol content. (A) Western blot analysis of cell lysates from control cells and NUDT15 and NUDT18 overexpressing cells, respectively, confirming NUDT15 and NUDT18 overexpression. (B) Schematic description of the setup of the cholesterol determination experiment using control and NUDT15/NUDT18 overexpressing cells. Figure was created with Biorender.com. (C) Relative total cholesterol content of NUDT15 overexpressing and control cells with and without mevastatin treatment. (D) Relative total cholesterol content of NUDT18 overexpressing and control cells with and without mevastatin treatment. Graphs shows the average and standard deviation from two independent experiments performed with two replicates.

We observed a statistically significant lower level of total cholesterol in NUDT15 overexpressing cells under untreated conditions compared to control cells. A similar trend, although not statistically significant, was observed in statin-treated cells (Figure 8C and Figure S6C). In the case of NUDT18 overexpression, we observed statistically significant lower cholesterol levels under both untreated and statin-treated conditions. Additional experiments consistently showed the same trend, although the effect on the total cholesterol content varied somewhat between experiments (Figure S6A and B). Taken together, these results imply that NUDT15 and NUDT18 catalyzed isoprene PP hydrolysis also occurs in the cell.

## DISCUSSION

### Catalysis of isoprene PP hydrolysis is a novel activity of NUDT15 and NUDT18

NUDT18 was initially identified as an enzyme capable of catalyzing the hydrolysis of 8-oxo-dGDP to 8-oxo-dGMP, suggesting a role in protecting against mutagenesis and DNA damage resulting from incorporation of 8-oxo-dG into DNA [25]. However, subsequent findings revealed that the depletion of NUDT18 neither increased the sensitivity to hydrogen peroxide, nor did overproduction of NUDT18 in MutT-deficient *E. coli* suppress the mutator phenotype [26]. This implies that NUDT18 may not primarily function in safeguarding the genome against oxidative stress but likely serves other cellular roles. NUDT15, closely related to NUDT1 (MutT homologue 1, MTH1) and commonly referred to as MTH2, was initially thought to act as a redundancy factor for MTH1 by catalyzing the hydrolysis of the mutagenic 8-oxo-dGTP [27]. However, NUDT15 was found to exhibit very weak 8-oxo-dGTP activity compared to MTH1 and no increase in 8-oxo-dG in DNA was observed upon NUDT15 depletion, questioning this view [22]. Recent studies have revealed significant roles for NUDT15 in the metabolism of anti-cancer drugs (thiopurines) and anti-viral drugs (aciclovir and ganciclovir) [28–31]. Nevertheless, the natural functions of NUDT15 have remained elusive.

Here, we present, for the first time, evidence that NUDT15 and NUDT18 exhibit pronounced activities with physiologically relevant isoprene PPs, suggesting a potential entirely novel cellular function for these enzymes. In comparison to previously reported activities of NUDT15 and NUDT18 with endogenous substrates, both enzymes demonstrate considerably higher catalytic efficiencies towards GPP. For NUDT18 the catalytic efficiency with GPP is approximately two times higher than the catalytic efficiency reported for 8-oxo-dGDP [21] and for NUDT15 the catalytic efficiency is more than 10 times higher than the corresponding value reported for this enzyme with 8-oxo-dGTP [32].

Figure 9 illustrates the isoprene biosynthesis pathway leading to production of cholesterol and other biomolecules and highlights the enzymes that are involved in isoprene PP metabolism in animals. The activities of NUDT18 and NUDT15 with isoprene PPs that we report here suggest that these enzymes have a role in modulating the cellular levels of these metabolites and, consequently their downstream products. Hydrolysis of GGPP and FPP, or other isoprene PPs used for their production, would decrease the concentration of available substrates for geranylgeranylation and farnesylation respectively. This reduction could lead to a decrease in the localization of small GTPases to cellular membranes, potentially affecting many cellular processes. Furthermore, GGOH and GOH are known key players in the feedback regulation of cholesterol [12, 33, 34]. They have also been shown to act as cellular signaling molecules and possess anti-inflammatory properties [35, 36]. Additionally, the isoprenoid alcohol Farnesol (FOH) has been identified as an activator of the nuclear farnesoid X hormone receptor [37]. These findings suggest that isoprene alcohols play roles beyond regulating isoprene biosynthesis. The source of GGOH, GOH and FOH are their corresponding isoprene pyrophosphates and enzyme catalyzed hydrolysis is likely required for their production. How these isoprenoid alcohols are produced in human cells is not fully understood, however, the polyisoprenoid diphosphate phosphatase (PDP1), integrated into the ER membrane, has been demonstrated to hydrolyze GGPP, FPP and GPP [18]. However, due to the membrane bound nature of this enzyme and the use of micelles in the activity experiment, a proper comparison of its activity with that of NUDT15 and NUDT18 with these isoprene PPs is not feasible. In contrast to PDP1, NUDT15 and NUDT18 are located in the soluble cytosolic fraction, like many other enzymes in the isoprenoid biosynthesis pathway such as Geranyl pyrophosphate synthase (GPPS) and Farnesyl pyrophosphate synthase (FPPS) (Figure 9) [38, 39]. Our results suggest that NUDT15 and NUDT18 may be responsible for the hydrolysis of isoprene PPs in the cytosol. Alternatively, these NUDIX enzymes may act in concert with PDP1 in the production of isoprenoid alcohols. An isoprenoid shunt, capable of converting isoprenoid alcohols into isoprene PPs via phosphorylation of isoprenoid alcohols and monophosphates, and vice versa, has been postulated to exist (Figure 9) [40]. The observed activities of NUDT15 and NUDT18 presented here imply a potential involvement of these enzymes in such a shunt.

**Figure 9.**
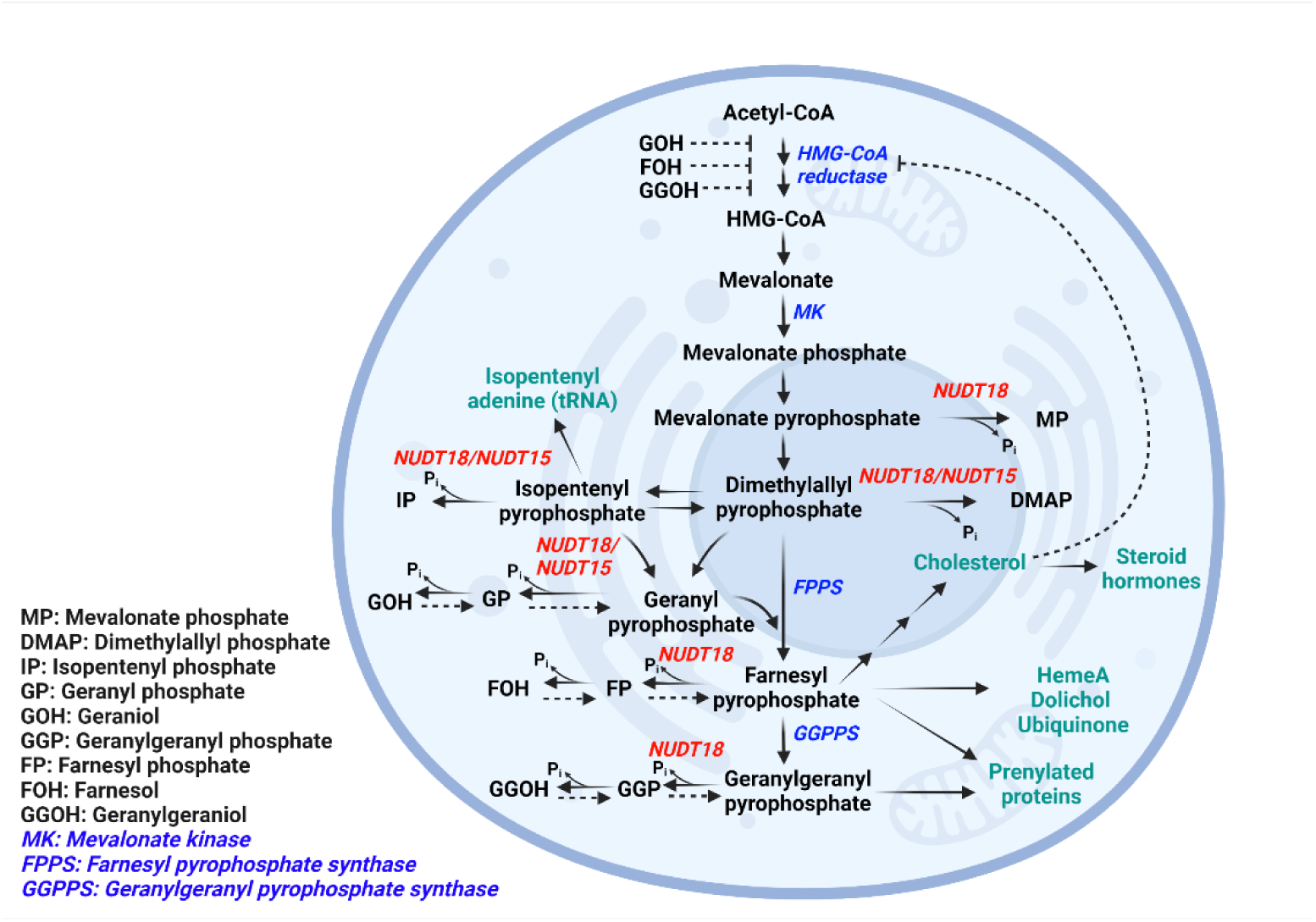
Schematic figure showing the biosynthetic pathways leading to cholesterol and other biomolecules (shown in dark cyan) where the isoprene pyrophosphates discovered to be substrates for NUDT18 are central. Enzymes discussed in the text are indicated in blue italics. NUDT15 and NUDT18 are indicated in red italics. Inhibition of HMG-CoA reductase catalyzed reaction by metabolites are indicated by dashed lines. Phosphorylation reactions are indicated by dashed arrows. Figure was created with Biorender.com.

### Relevance of NUDIX enzyme catalyzed isoprene PP hydrolysis in the cell

Does the hydrolysis activity of NUDT15 and NUDT18 with isoprene PPs that we observe in our biochemical activity assay *in vitro* also manifest in a cellular context? The catalytic efficiency of NUDT18 with GPP was determined to be 2.1·10^5^ M^-1^s^-1^ at 22 °C, a value comparable to the reported catalytic efficiency determined at 37 °C for geranylgeranyl pyrophosphate synthase (GGPPS) and FPPS [41, 42], enzymes catalyzing the formation of GGPP and FPP in the cholesterol synthesis pathway, respectively (Figure 9). The catalytic efficiency of NUDT18 at 37 °C is expected to be considerably higher than the catalytic efficiency determined at 22 °C. This suggests that NUDT18 is likely to hydrolyze GPP within the cell. Furthermore, the median kcat/Km value from a dataset comprising published values for approximately 1,900 enzymes with their natural substrates is around 1·10^5^ [43]. The two-fold higher catalytic efficiency of NUDT18 with GPP reported here supports the likelihood that GPP is an endogenous substrate of NUDT18. Additionally, the low Km value of NUDT18 for GPP (3.7 µM) (Table 1) is comparable to the Km value of geranylgeranyl pyrophosphate synthase (GGPPS) for IPP and FPP [41, 42], as well as the Km values of Farnesyl pyrophosphate synthase (FPPS) for IPP and GPP [42]. This further supports the conception that NUDT18-catalyzed hydrolysis of GPP occurs within the cell. Cellular hydrolysis of isoprene PPs, which constitute metabolites in the cholesterol synthesis pathway, would ultimately impact cholesterol production since FPP, which is utilized for cholesterol production, is synthesized from GPP and DMAPP (Figure 9). Consequently, we hypothesized that overexpression of NUDT18 and NUDT15 would lead to a decrease in cholesterol levels due to increased hydrolysis of these metabolites, even though activity with FPP is low. Indeed, our findings indicate that the overexpression of NUDT15 leads to a reduction in cholesterol levels in untreated cells. Furthermore, the overexpression of NUDT18 results in a decrease in total cholesterol levels in both untreated and statin-treated cells, consistent with the suggested involvement of these enzymes in isoprene PP metabolism. However, further studies are warranted to investigate the specific impact of NUDT15 and NUDT18 activities on the cellular levels of individual isoprene PP metabolites.

Both NUDT15 and NUDT18 exhibit low tissue specificity, being expressed in most tissues (Human Protein Atlas, https://www.proteinatlas.org). Elevated cellular levels of NUDT15 and NUDT18 could reduce cholesterol production by decreasing the concentration of intermediates in the cholesterol synthesis pathway (Figure 9). Interestingly, in the liver, where cholesterol production is high, the level of NUDT18 mRNA is very low and the protein is undetectable (Human Protein Atlas, https://www.proteinatlas.org). Moreover, cells defficient in NUDT18 display a significantly slower proliferation rate, suggesting a role of NUDT18 in cell growth [26]. We speculate that this effect may stem from NUDT18 modulating the concentration of substrates available for protein isoprenylation or other isoprene metabolites. We anticipate that our findings will inspire investigations into isoprene dependent metabolites in clinical samples from patients carrying *nudt15* gene variants, resulting in low NUDT15 enzyme levels and consequently reduced cellular NUDT15 activity [30, 44, 45].

In conclusion, we here present a novel and significant activity of human NUDT15 and NUDT18 in catalyzing the hydrolysis of isoprene PPs, and propose that these enzymes could play a role in isoprenoid metabolism in animals.

## EXPERIMENTAL PROCEDURES

### Protein production

The human NUDIX enzymes MTH1 (NUDT1), NUDT2, NUDT3, NUDT4, NUDT7, NUDT9, NUDT10, NUDT11, NUDT12, NUDT14, NUDT15, NUDT16, NUDT17, NUDT18, NUDT21 and NUDT22 were produced as described previously [46]. In brief, NUDIX enzymes were expressed in *E. coli* as N-terminally His-tagged proteins and purified using His-Trap HP followed by gel filtration or cation exchange chromatography. NUDT15His49Ala cDNA was purchased from GeneArt and subcloned into pNIC28 (a kind gift from PSF, Karolinska Institutet) using NdeI and SalI. NUDT15His49A was expressed in BL21 (DE3) T1R pRARE2 at 18 °C overnight after induction with 0.5 mM IPTG. The sonicated lysate was centrifuged, filtered, and purified using His-Trap HP (GE Healthcare) followed by gel filtration using HiLoad 16/60 Superdex 75 columns on an ÄKTA Xpress. The purity of purified proteins was assessed using SDS-PAGE analysis. Proteins were stored in storage buffer (20 mM HEPES, 300 mM NaCl, 10% glycerol, 2 mM TCEP, pH 7.5) and kept as aliquots at -80 °C.

### Screening of NUDIX enzymes for activity with isoprene PPs

Hydrolysis activity of a panel of 17 human NUDIX proteins with DMAPP and IPPP was tested by incubating 100 µM of each isoprene PP with 100 nM human NUDIX enzyme for 30 minutes at 22°C. Formed Pi was detected by addition of Malachite green reagent [47] followed by measurement of absorbance at 630 nm, after incubation at 22 °C for 15 minutes. Activity of NUDT15 and NUDT18 (100 nM) was also tested with 100 µM Mevalonate pyrophosphate (MPP, Sigma-Aldrich, #94259), IPPP (Sigma-Aldrich, I0503), DMAPP (Cayman Chemicals and Sigma-Aldrich D4287), Geranyl-PP (GPP, Sigma-Aldrich, G6772), Farnesyl-PP, (FPP, Sigma-Aldrich, F6892) and GGPP (Sigma G6025) in assay buffer (100 mM Tris acetate pH 8.0, 40 mM NaCl, 10 mM Magnesium acetate and 0.01% Tween 20) and compared to the activity with 100 µM of the previously identified substrates dGTP (GE Healthcare, 27-1870-04) for NUDT15 and 8-oxo-dGDP (Jena Bioscience, NU-1158) for NUDT18. In addition, since NUDT15 is closely related to MutT homolog 1 (MTH1, NUDT1) the activity of 50 nM *E. coli* MutT with 100 µM GPP as well as with 100 µM dGTP as a control was tested as described above.

### LC-MS analysis of NUDT15 and NUDT18 catalyzed GPP and FPP hydrolysis products

1 mM GPP was incubated with 0.5 µM NUDT18 or 1 µM NUDT15 in reaction buffer (0.1 M Tris acetate pH 8.0, 40 mM NaCl, 10 mM Magnesium acetate) at 22 °C with rotation. The reaction was stopped at different time points, ranging from 0 to 180 minutes for NUDT18 and 0 to 120 minutes for NUDT15, by removing an aliquot to which EDTA was added (90 mM final concentration) in order to chelate Mg^2+^ ions needed for catalysis. The samples were then heated for 5 minutes at 75 °C to denature the enzyme, followed by centrifugation at 17,000 *g* for 10 minutes to precipitate the denatured protein. The supernatants were analyzed using an Agilent 1100 HPLC system equipped with an X-bridge C18 column 5 µm (3x 50 mm) coupled to an Agilent MSD mass spectrometer. For analysis of FPP hydrolysis the same procedure was used with the modification that 0.5 mM FPP was incubated with 10 µM NUDT15 or NUDT18 and samples were taken at various time points between 0 and 225 minutes, and then prepared and analyzed as described above for the analysis of GPP hydrolysis.

### Detailed kinetic analysis of NUDT15 and NUDT18 with isoprene PPs

To further analyze the activity of NUDT18 and NUDT15 with isoprene PPs a detailed kinetic characterization was performed with a panel of isoprene PPs. Initial rates were determined using 25 nM of NUDT18 and substrate concentrations ranging from: 0-50 µM for GPP and 0-100 µM for MPP, DMAPP, IPPP, FPP and GGPP. For saturation curves produced with NUDT15 25 nM enzyme was used and substrate concentration ranges used were: 0-50 µM for GPP, 0-200 µM for DMAPP and 0-150 µM for IPPP. Activities were assayed in reaction buffer (100 mM Tris acetate pH 8.0, 40 mM NaCl, 10 mM Magnesium acetate and 0.01% Tween 20). Reactions were stopped after 10, 20 and 30 minutes incubation time at 22 °C and inorganic phosphate (Pi), released upon hydrolysis, was detected by addition of malachite green reagent and measurement of the absorbance at 630 nm using a Hidex plate reader. A Pi standard curve on the assay plate was used to convert absorbance to amount of Pi produced. Initial rates were determined in duplicate and saturation curve experiments were performed twice apart from for NUDT15 activity with FPP and GGPP which only was performed once due to low activity. Kinetic parameters were determined by fitting the Michaelis-Menten equation to initial rate data using GraphPad Prism 8.0.

### Activity analysis of NUDT15wt, NUDT15H49A and MutT with dGTP and GPP

Activity with 100 µM GPP was assayed in assay buffer (0.1 M Tris acetate pH 8.0, 40 mM NaCl, 10 mM Magnesium acetate and 0.01% Tween20) using 100 nM NUDT15H49A, NUDT15WT or MutT. Activity with 100 µM dGTP was monitored in assay buffer fortified with PPase 0.4U/ml, converting PPi to Pi, and 50 nM NUDT15H49A, 5 nM NUDT15WT or 50 nM MutT. Formed Pi was detected after incubation for 30 minutes at 22 °C by addition of Malachite green reagent. A Pi standard curve was used to calculate the concentration of produced Pi. Saturation curves of NUDT15WT and NUDT15H49A were produced by determining initial rates in assay buffer (0.1 M Tris acetate pH 8.0, 40 mM NaCl, 10 mM Magnesium acetate) using 50 nM NUDT15 WT or 100 nM NUDT15H49A and substrate concentrations of GPP ranging from 0 to 200 µM. For saturation curves with dGTP, dGTP was varied between 0 and 400 µM and initial rates of 5 nM NUDT15WT and 50 nM NUDT15H49A were monitored.

### Crystallization and structure determination of the NUDT15-GP complex

Purified human NUDT15 (20 mg/mL) was preincubated with 10 mM GPP (Sigma-Aldrich) for 2 hours at 20 °C. The protein was crystallized using sitting drop vapor diffusion at 20 °C in 0.1 M Tris-HCl pH 8.5, 0.2 M sodium acetate and 30 % PEG4000. Protein crystals were flash frozen directly in liquid nitrogen without adding additional cryoprotectant. X-ray diffraction data was collected at station I03 of the Diamond Light Source (Oxford, UK) equipped with a PILATUS-6M detector. A complete data set was collected on a single crystal at 100 K. The dataset was processed and scaled with DIALS [48] and Aimless [49] within the CCP4 suite [50]. Molecular replacement was performed in Phaser [51], using a human NUDT15 structure (PDB ID: 5LPG) with all ligands and waters removed. Several rounds of manual model building and refinement were performed using COOT [52] and REFMAC5 [53] during which waters and the ligand GP were added to the structure. Data processing and refinement statistics are listed in Table S1. The coordinates and structure factors for the NUDT15-GP structure presented in this paper were deposited in the PDB under the accession code 7R0D.

### Generation of NUDT15 and NUDT18 overexpressing stable cell lines

Codon-optimized NUDT15 (NM_018283.3) and NUDT18 (NM_024815.3) cDNA including a Kozak sequence (TCCACCATGG) at the 5’-end were obtained as gene fragments (Integrated DNA Technologies). The NUDT15 and NUDT18 cDNA were subcloned into the entry vectors, N-3xFlag-pENTR4 and pENTR4-N-3HA, respectively, using SalI and NotI restriction sites. Making use of the Gateway cloning technique, the cDNA encoding the HA-tagged NUDT18 and the Flag-tagged NUDT15 was transferred into the destination vectors pLenti-PGK-Hygro DEST (addgene #19066) and pLenti-PGK-Blast DEST (addgene #19065), respectively. Stable lentiviruses carrying tagged-NUDT15 or NUDT18 cDNA were produced by transfecting HEK293T with a mixture of vector DNA and packaging plasmids (addgene, #12260 and #12259). Virions were harvested after 36 and 60 hours post transfection. Breast cancer cell line BT549 was obtained from ATCC and was cultured in RPMI 1640 medium supplemented with 10% fetal bovine serum, at 37 °C in 5% CO_2_ humidified atmosphere. Virions were transduced into BT549 breast carcinoma cells and stably expressing cell lines were produced by growing the cells in RPMI 1640 (Thermo Fisher #61870044) + 5 % FBS (Gibco #10500064) under antibiotic pressure using 5 μg/ml blasticidin (Gibco #A1113903) for HA-NUDT18 overexpressing and HA-V control cells and 200 μg/ml hygromycin B (Gibco #10687010) for Flag-NUDT15 overexpressing and Flag-V control cells. Overexpression of Flag-NUDT15 and HA-NUDT18 were confirmed by analyzing cell lysates by SDS-PAGE followed by Western blotting using anti-FLAG-tag Rabbit antibody (#740001, Thermo Fischer Scientific) and anti-HA-tag (6E2) Mouse antibody (#2367, Cell Signaling) followed by anti-Rabbit HRP or anti-Mouse HRP. Protein bands were detected using the SuperSignal West Femto Maximum Sensitivity Substrate (**#**34096, Thermo Scientific). Images were collected using an Odyssey Fc Li-COR imager.

### Determination of cholesterol content in NUDT15 and NUDT18 overexpressing cells

Cells were seeded in T75 flasks in OptiPRO (Gibco #12309019) + 1% dialyzed FBS (Gibco #A3382001), one flask for each cell line and condition. After culturing for 24 hours, mevastatin (5 µM, Sigma-Aldrich #M2537), or the corresponding volume of solvent, was added to the cell cultures. Before its addition, mevastatin was activated by neutralization with an equal amount of 1 M HCl followed by dilution to 10 mM with cell medium + 50% EtOH. Cells were cultivated for an additional 48 hours after which the cells were washed once with DPBS (Gibco #14190169), trypsinized, detached and resuspended in seeding medium. Cells were washed once in ice-cold DPBS and resuspended in 1 ml DPBS and transferred to Eppendorf tubes. Cell counts were measured in triplicate using a TC20 Automated cell counter (BioRad) (15 µl cell suspension + 15 µl Trypan Blue Dye (0.4% solution, BioRad #1450013), 10 µl was transferred to each chamber in a cell counting slide with dual chambers (BioRad, #145-0011)). The lipid fraction from 1 million cells per sample was extracted after which total cholesterol was assayed using the Cholesterol/Cholesteryl Ester Quantitation Assay kit (Abcam, ab65359) according to the manufacturer’s recommendations. Experiments were performed at least twice under identical conditions and cholesterol content was assayed in duplicate. An additional experiment analyzing the effect of NUDT15 overexpression was performed using 2.5 µM mevastatin. Statistical significance was calculated using TwoWay Anova and GraphPad Prism 8.0.

## Data availability

All data can be shared upon request made to the corresponding author. Crystal structure data is deposited under the PDB under the accession code 7R0D.

## Supporting information

This article contains supporting information.

## Acknowledgments

We thank Athina Pliakou, Louise Sjöholm, Kristina Edfeldt, Mari Kullman Magnusson, Flor Pineiro and Sabina Eriksson for their assistance in the Helleday Lab. We thank the beamline scientists at Diamond Light Source (United Kingdom, proposal mx21625) for their support in structural biology data collection. We are grateful to the Protein Science Facility at Karolinska Institutet/SciLifeLab (http://ki.se/psf) for help with protein production.

## Author Contributions

A.S.J wrote the manuscript with input from E.S. and P.S.. All authors commented on the manuscript. A.S.J. and P.S. conceived the project. A.S.J. designed, performed, and analysed all biochemical experiments. D.K. performed western blotting experiments. J.U. prepared the stably overexpressing cell lines. T.K. and K.V. performed the LC-MS analysis. A.S.J. and I.A. designed and executed cell biology experiments. Protein structure was solved by E.S. and P.S. and E.S performed structural analyses. A.S.J., P.S. and T.H. supervised the project and acquired funding.

## Funding and additional information

This work was supported by grants from the Swedish Research Council (2022-03681 to P.S., 2015-00162 and 2017-06095 to T.H.), the Swedish Cancer Society (20 1287 PjF to P.S. and 21 1490 Pj to T.H.), the European Research Council (ERC-AdG-TAROX-695376 to T.H.), the Knut and Alice Wallenberg Foundation (KAW2014.0273 to T.H.), Torsten and Ragnar Söderberg Foundation (to T.H.), the Helleday Foundation (to J.U.) and KI funds (2020-02211 to A.S.J).

## Abbreviations

The abbreviations used are: NUDIX, <u>nu</u>cleoside <u>di</u>phosphates linked to a number of different moieties (<u>x</u>); IPPP, isopentenyl pyrophosphate; DMAPP, dimethylallyl pyrophosphate, TCEP, tris(2-carboxyethyl)phosphine; MK, Mevalonate kinase; GPP, geranyl pyrophosphate; GP, Geranyl monophosphate; FPP, farnesyl pyrophosphate; FP, farnesyl monophosphate; GGPP, geranylgeranyl pyrophosphate; HMG-CoA reductase, 3-hydroxy-3-methylglutaryl-CoA reductase; GGOH, geranylgeraniol; GOH, geraniol; FOH, farnesol; NUDT15, NUDIX hydrolase 15; NUDT18, NUDIX hydrolase 18; PP, pyrophosphate; AtNUDT1, *Arabidopsis thaliana* NUDIX hydrolase 1; GGPPS, geranylgeranyl pyrophosphate synthase; FPPS, farnesyl pyrophosphate synthase; 8-oxo-dGDP, 8-Oxo-2’-deoxyguanosine-5’-diphosphate; dGTP, deoxyguanosine-5’-triphosphate; PPase, pyrophosphatase, PPi, inorganic pyrophosphate; Pi, inorganic phosphate; MPP, Mevalonate pyrophosphate; FBS, Fetal bovine serum; DPBS, Dulbecco′s Phosphate Buffered Saline; IPTG, Isopropyl β-D-1-thiogalactopyranoside

