## Supplementary information for "Kinetic and structural characterization of human NUDIX hydrolases NUDT15 and NUDT18 as catalysts of isoprene pyrophosphate hydrolysis"

**This PDF file includes:**

Figures S1 to S6

Table S1-S2

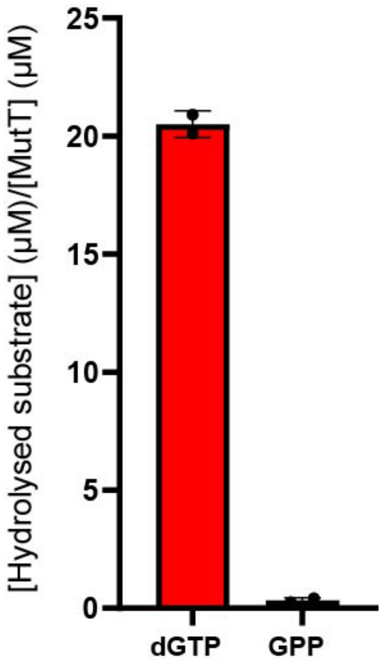


**Figure S1. MutT does not display any activity with GPP.** *E. coli* MutT, a bacterial homologue of MTH1 and NUDT15, was tested for activity with GPP (100 µM) and with dGTP (100 µM) as a positive control using 50 nM enzyme. Graph shows results from one experiment with data points in duplicate.

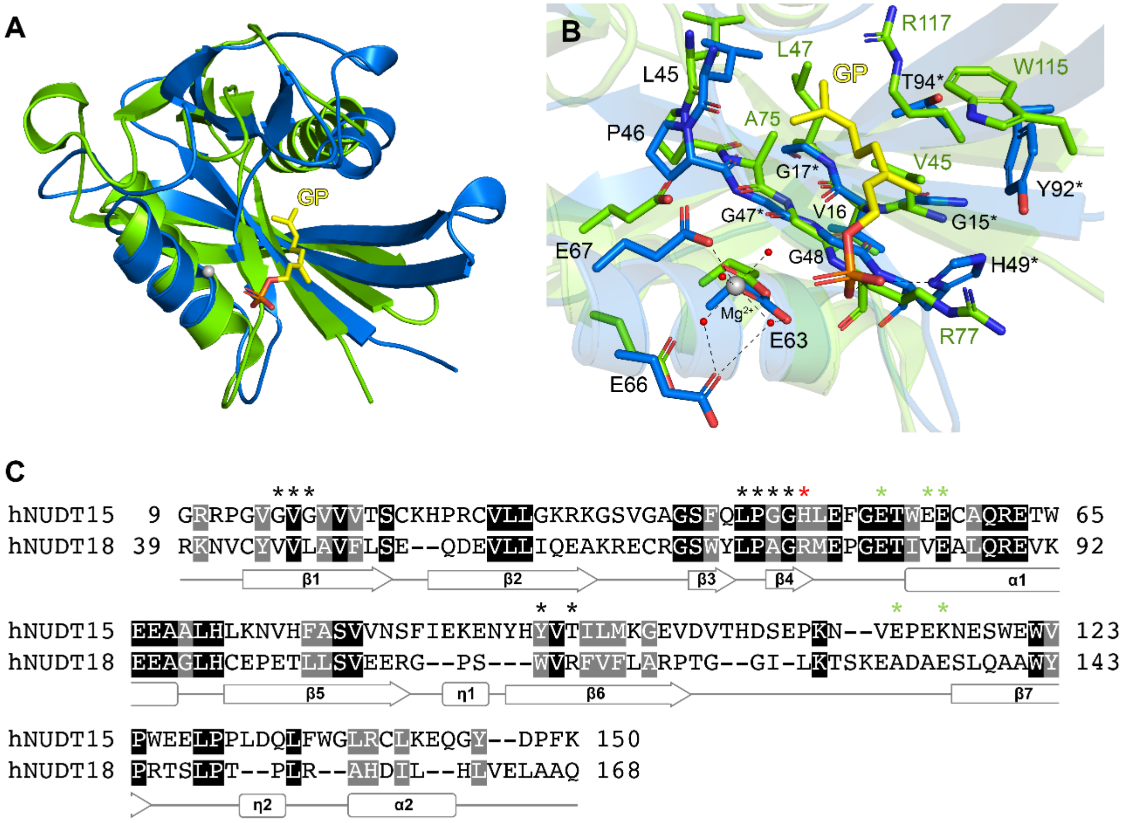


**Figure S2. Structural comparison of hNUDT15-GP with the hNUDT18 nucleotidase domain.** (A) Superposition of hNUDT15-GP (blue) and AtNUDT1 (green, PDB ID: 3GG6) monomers. GP is shown as a yellow stick model with the coordinated magnesium ion shown as a grey sphere. (B) Comparison of the residues involved the coordination of GP and magnesium in the hNUDT15-GP structure. Residue numbering represents the hNUDT15 structure and in cases where the residue is not conserved, the residue from hNUDT15 is labelled with an asterisk and the corresponding residue from AtNUDT1 is labelled in dark green. Nitrogen, oxygen and phosphor atoms are coloured blue, red and orange, respectively. Water molecules are shown as red spheres. Hydrogen bond interactions are shown as dashed lines. (C) Amino acid sequence alignment of human NUDT15 (UniProt: Q9CA40) and the nucleotidase domain of human NUDT18 (Q6ZVK8) performed using Clustal Omega through the EBI webserver. The resulting alignment is coloured according to sequence similarity using BOXSHADE. Identical residues are shaded black, while grey shading indicates amino acids with conserved physicochemical properties. Residues in hNUDT15 which form hydrogen bond or hydrophobic interactions with GP are shown as asterisks, coloured red and black, respectively. Green asterisks indicate residues required for magnesium coordination. The secondary structure corresponding to the amino acid sequence of hNUDT15-GP is displayed below the alignment.

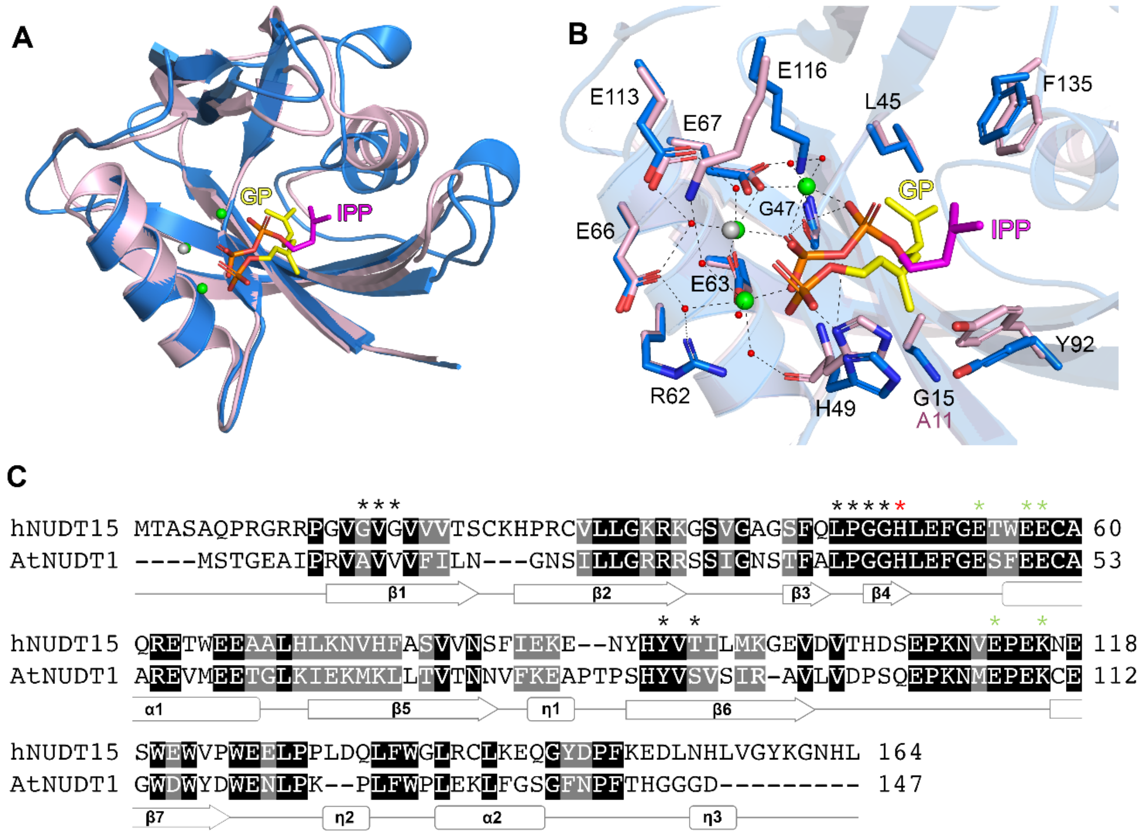


**Figure S3. Structural comparison of hNUDT15-GP with AtNUDT1-IPP.** (A) Superposition of hNUDT15-GP (blue) and AtNUDT1 (light pink, PDB ID: 6DBZ) monomers. GP and IPP are shown as sticks, coloured yellow and magenta, respectively. Magnesium ions are coloured grey (hNUDT15) or green (AtNUDT1). (B) Comparison of the residues involved the coordination of IPP and magnesium in the AtNUDT1-IPP structure. Residue numbering represents the hNUDT15 structure and in cases where the residue is not conserved, the relevant residue from AtNUDT1 is labelled in dark pink. Nitrogen, oxygen and phosphor atoms are coloured blue, red and orange, respectively. Water molecules are shown as red spheres. Hydrogen bond interactions are shown as dashed lines. (C) Amino acid sequence alignment of human NUDT15 (UniProt: Q9CA40) and *A. thaliana* NUDT1 (Q9NV35) performed using Clustal Omega through the EBI webserver. The resulting alignment is coloured according to sequence similarity using BOXSHADE. Identical residues are shaded black, while grey shading indicates amino acids with conserved physicochemical properties. Residues in hNUDT15 which form hydrogen bond or hydrophobic interactions with GP are shown as asterisks, coloured red and black, respectively. Green asterisks indicate residues required for magnesium coordination. The secondary structure corresponding to the amino acid sequence of hNUDT15-GP is displayed below the alignment.

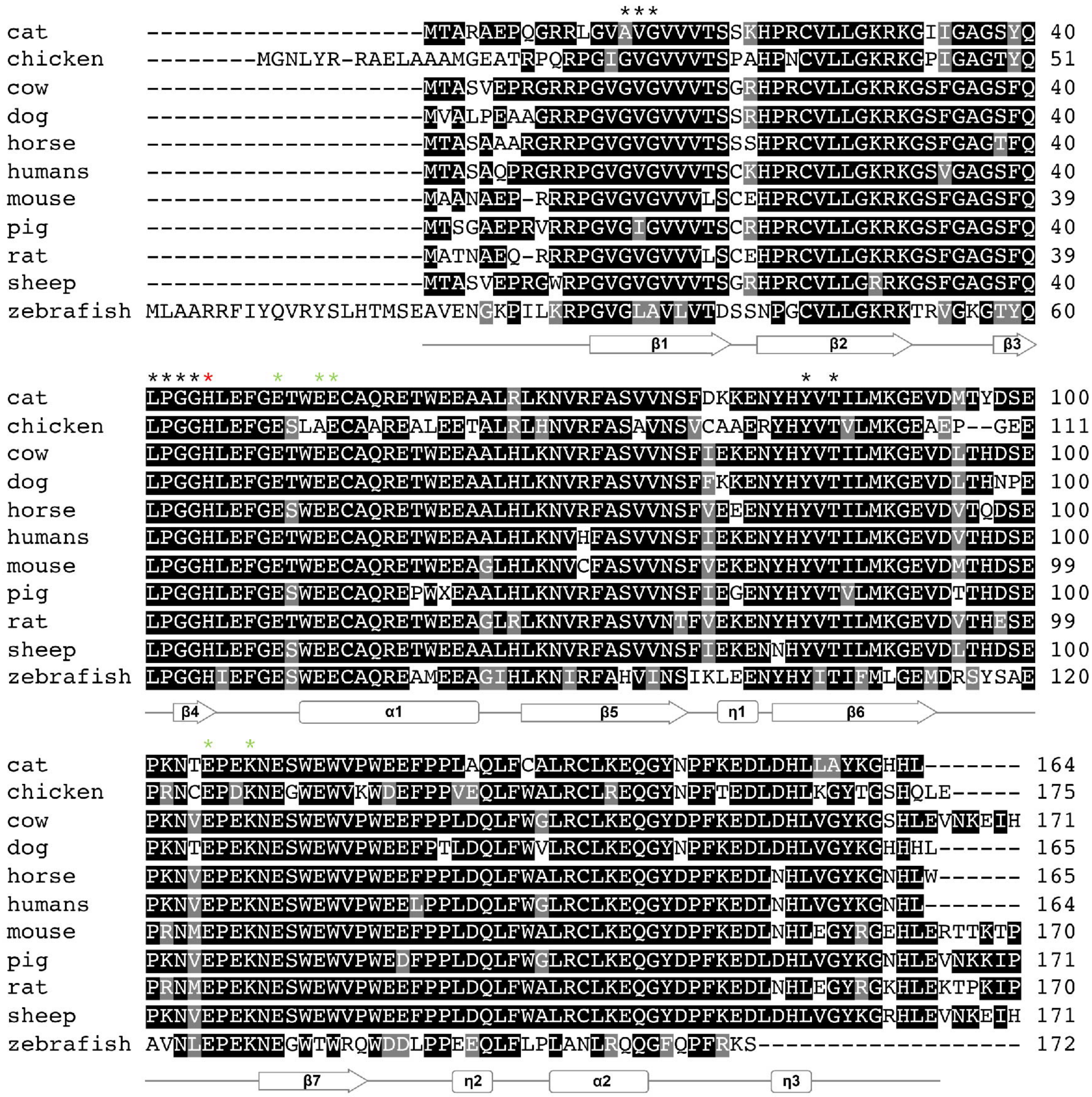


**Figure S4. Sequence alignment of human and mammalian NUDT15 proteins.** NUDT15 amino acid sequences from *F. catus* (UniProt: A0A2I2U059), *G. gallus* (UniProt: F1NV92), *B. taurus* (UniProt: E1B7T3) *C. lupus* (UniProt: ER2DP5), *E. caballus* (UniProt: F7BQG1), *H. sapiens* (UniProt: Q9NV35), *M. musculus* (UniProt: Q8BG93), *S. scrofa* (UniProt: I3LBG2), *R. norvegicus* (UniProt: D3ZKQ0), *O. aries* (UniProt: A0A6P3THI6) and *D. rerio* (UniProt: Q6IQB3) were compared using Clustal Omega through the EBI webserver. The resulting alignment is coloured according to sequence similarity using BOXSHADE. Identical residues are shaded black, while grey shading indicates amino acids with conserved physicochemical properties. Residues from the hNUDT15-GP structure shown to form hydrogen bonds or hydrophobic interactions with GP are shown as asterisks, coloured red and black, respectively. Green asterisks indicate residues required for magnesium coordination. The secondary structure corresponding to the amino acid sequence of hNUDT15-GP is displayed below the alignment.

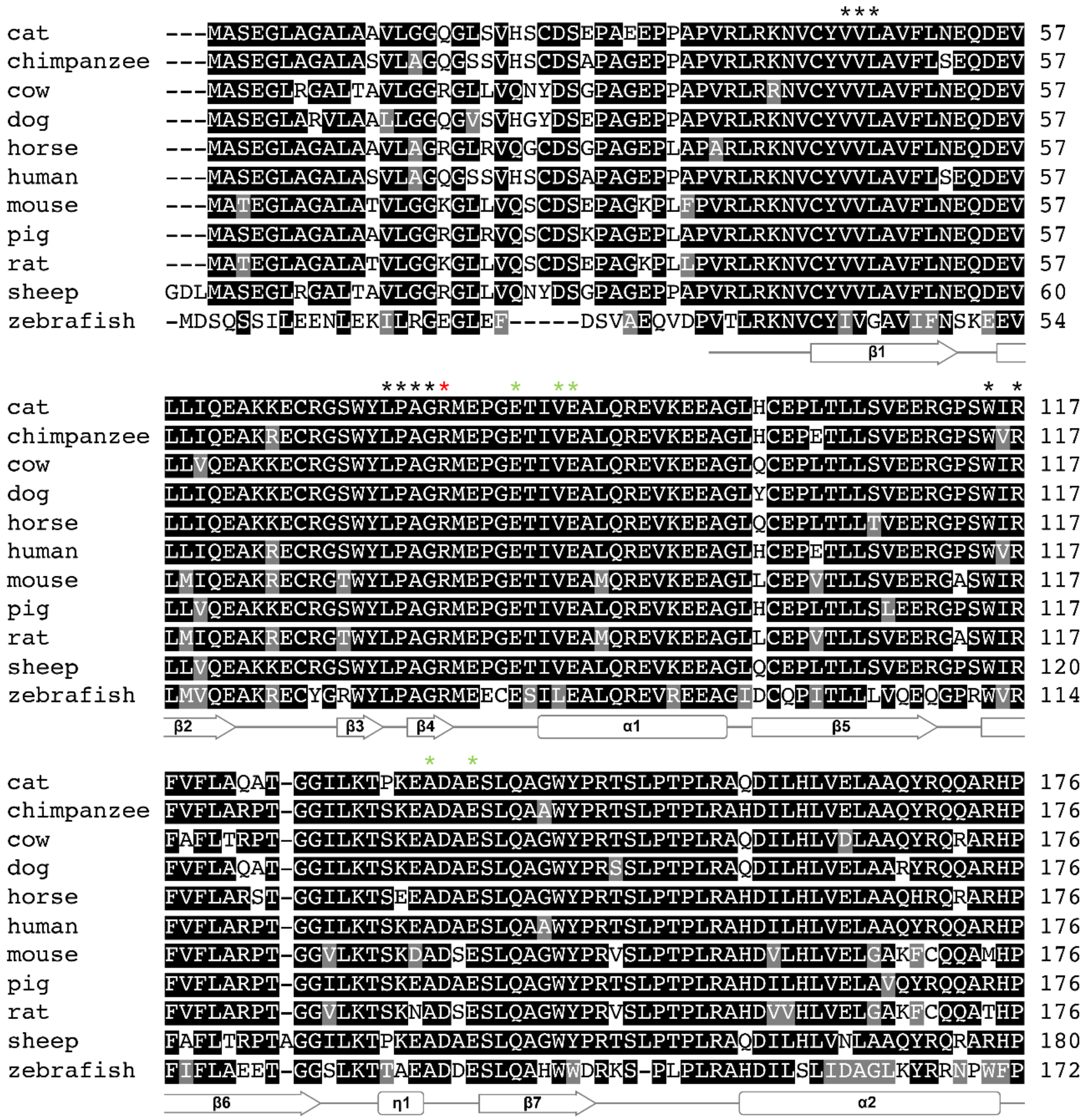


**Figure S5. Sequence alignment of the NUDT18 nucleotidase domain from humans and other species.** NUDT18 nucleotidase amino acid sequences from *F. catus* (NCBI: XP_003984764.1), *P. troglodytes* (UniProt: H2QVU3), *B. taurus* (UniProt: F1N0N5) *C. lupus* (UniProt: F1PDW5), *E. caballus* (UniProt: F6RHM5), *H. sapiens* (UniProt: Q6ZVK8), *M. musculus* (UniProt: Q3U2V3), *S. scrofa* (UniProt: F1RMB9), *R. norvegicus* (UniProt: Q641Y7), *O. aries* (UniProt: W5PLD7) and *D. rerio* (UniProt: Q568Q0) were compared using Clustal Omega through the EBI webserver. The resulting alignment is coloured according to sequence similarity using BOXSHADE. Identical residues are shaded black, while grey shading indicates amino acids with conserved physicochemical properties. Residues from human NUDT18 predicted to interact with isoprenoids (specifically GP) are shown as asterisks, coloured red and black, respectively. Green asterisks indicate residues required for magnesium coordination. The secondary structure corresponding to the amino acid sequence of the hNUDT18 nucleotidase domain (PDB ID: 3gg6) is displayed below the alignment.

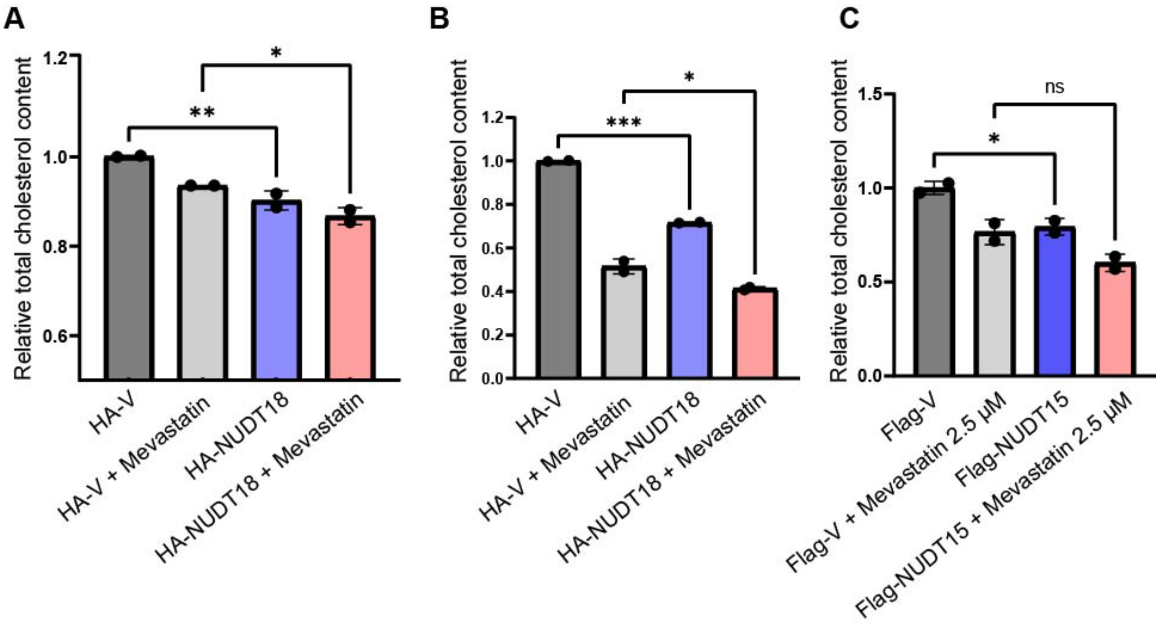


**Figure S6. Overexpression of NUDT15 and NUDT18 decreases total cholesterol content.** (A) and (B) Control cells (HA-V) and NUDT18 overexpressing cells (HA-NUDT18) were treated with 5 µM mevastatin or left untreated 24 hours after seeding. Cells were harvested after an additional 48 hours of culturing. Lipid fraction was extracted and cholesterol content was determined. (C) Control cells (Flag-V) and NUDT15 overexpressing cells (Flag-NUDT15) were treated as in A and B but with 2.5 µM mevastatin, the lipid fraction was extracted and cholesterol content was determined. Graphs show independent experiments and present the average and standard deviation of samples assayed in duplicate.

**Table S1.** Data Collection and Refinement Statistics

| | NUDT15-GP |
|---|---|
| PDB code | 7R0D |
| **Data collection** | |
| Space group | $P2_12_12_1$ |
| Cell dimensions | |
| *a, b, c* (Å) | 46.7, 48.3, 135.2 |
| $\alpha, \beta, \gamma$ (°) | 90.0, 90.0, 90.0 |
| Resolution (Å) | 46.6-1.70 (1.73-1.70) |
| No. observations | 388242 (8976) |
| No. unique reflections | 34417 (1759) |
| $R_{merge}$ | 0.07 (0.53) |
| CC(1/2) | 0.99 (0.90) |
| $I/\sigma I$ | 17.8 (2.6) |
| Completeness (%) | 99.7 (96.7) |
| Redundancy | 11.3 (5.1) |
| **Refinement** | |
| Resolution (Å) | 45.5-1.70 |
| No. reflections | 32621 |
| $R_{work}/R_{free}$ (%) | 18.4/22.6 |
| No. of atoms | |
| Protein | 2974 |
| Ligand | 30 |
| Metal | 2 |
| Water | 172 |
| *B*-factors | |
| Protein (Å$^2$) | 29.4 |
| Ligand/ion (Å$^2$) | 52.3 |
| Metal (Å$^2$) | 34.7 |
| Water (Å$^2$) | 35.7 |
| RMSDs | |
| Bond lengths (Å) | 0.012 |
| Bond angles (°) | 1.71 |

Values in parentheses are for the highest-resolution shell

**Table S2. Kinetic parameters for NUDT15WT and NUDT15H49A with GPP and dGTP**

| | GPP | | | dGTP | | |
|---|---|---|---|---|---|---|
| | $k_{cat}$ ($min^{-1}$) | $K_M$ (µM) | $k_{cat}/K_M$ ($M^{-1}min^{-1}$) | $k_{cat}$ ($min^{-1}$) | $K_M$ (µM) | $k_{cat}/K_M$ ($M^{-1}min^{-1}$) |
| NUDT15WT | 8.9 | 31.7 | 280,420 | 38.2 | 21 | 1820,300 |
| NUDT15H49A | 0.3 | - | - | 19.0 | 289 | 65,780 |
